# The impact of real-time fMRI denoising on online evaluation of brain activity and functional connectivity

**DOI:** 10.1101/2021.03.02.433573

**Authors:** Masaya Misaki, Jerzy Bodurka

## Abstract

**Objective:** Comprehensive denoising is imperative in fMRI analysis to reliably evaluate neural activity from the blood oxygenation level dependent signal. In real-time fMRI, however, only a minimal denoising process has been applied and the impact of insufficient denoising on online brain activity estimation has not been assessed comprehensively. This study evaluated the noise reduction performance of online fMRI processes in a real-time estimation of regional brain activity and functional connectivity.

**Approach:** We performed a series of real-time processing simulations of online fMRI processing, including slice-timing correction, motion correction, spatial smoothing, signal scaling, and noise regression with high-pass filtering, motion parameters, motion derivatives, global signal, white matter/ventricle average signals, and physiological noise models with image-based retrospective correction of physiological motion effects (RETROICOR) and respiration volume per time (RVT).

**Main results:** All the processing was completed in less than 400 ms for whole-brain voxels. Most processing had a benefit for noise reduction except for RVT that did not work due to the limitation of the online peak detection. The global signal regression, white matter/ventricle signal regression, and RETORICOR had a distinctive noise reduction effect, depending on the target signal, and could not substitute for each other. Global signal regression could eliminate the noise-associated bias in the mean dynamic functional connectivity across time.

**Significance:** The results indicate that extensive real-time denoising is possible and highly recommended for real-time fMRI applications.

## 1 Introduction

Real-time functional magnetic resonance imaging (rtfMRI) is a system for evaluating online brain activity as soon as an imaging volume is acquired [1, 2]. Applications of rtfMRI have expanded from online inspection of signal quality [1] to brain-computer interface for manipulating equipment with brain signal [3] and, most significantly, to neurofeedback for self-regulating brain activity [4, 5]. The primary challenge of rtfMRI is performing image processing in a short time. The processing includes image reconstruction, data processing for denoising and extracting the target signal, and displaying the processed signal. All these processes need to be done in less than the repetition time (TR) of volume acquisition, usually a few seconds or less than 1 s, to keep the pace of real-time data presentation without accumulating a delay. Thanks to the advancement of computational hardware, image reconstruction and signal presentation times are becoming negligibly short. In contrast, the time of data processing for signal denoising is still not negligible and its impact on signal quality has not yet been well-examined [6].

The blood-oxygen-level-dependent (BOLD) signal in fMRI contains various noise and requires many denoising steps to get a reliable estimate of neural activity. Although comprehensive denoising is critical to obtain a reliable estimate of brain activity, its execution in a rtfMRI environment is so cumbersome that most rtfMRI applications have used minimal denoising steps compared to offline analyses. The real-time estimate of neural activity with a limited denoising process cannot have comparable quality to that available in offline processing. This could hinder the reproducibility of the result of rtfMRI applications like neurofeedback [7].

This limitation has been being compensated for through the implementation of a fast computation algorithm, the development of a processing method adapted for real-time computation, and the improvement of the computational capacity of personal computers and graphics processing units (GPU) [8–10]. Several packaged frameworks implementing advanced real-time processing have been released as well [11–13]. These improvements in real-time signal processing for fMRI have been reducing the gap between offline and real-time evaluations of neural activity, which should improve the efficacy of rtfMRI applications [14].

Notwithstanding that these advanced online processing systems could boost the reliability of neural activity estimates in rtfMRI, their use has not been considered obligatory in rtfMRI applications. Moreover, the impact of real-time denoising steps on the quality of the neurofeedback signal has not been well-recognized, demonstrated by the fact that most neurofeedback studies do not report real-time denoising steps in detail [6]. The CRED-nf checklist [15] is one of the efforts to resolve this undesired state of the field by encouraging researchers to report details of the real-time processing steps, while it does not indicate what processing is recommended for which rtfMRI application.

We have not known which type of real-time processing has a benefit in which rtfMRI application due to the dearth of studies quantifying the denoising benefit in improving neurofeedback signal quality. While a few reports are available about the benefit of fMRI real-time processing [8, 16–18], these reports focus on specific parts of the process and do not cover comprehensive denoising steps as employed in offline analysis. Heunis, Lamerichs, Zinger, Caballero-Gaudes, Jansen, Aldenkamp and Breeuwer [6] has described this state of research and highlighted the need for a methodological study that quantifies the effect of the denoising steps on rtfMRI signal quality.

This study aimed to comprehensively evaluate the benefit of fMRI real-time processing (RTP) in improving the quality of online neural activity estimates. Specifically, we performed RTP simulation analyses to evaluate how much each stage of RTP reduced the noise effect on the online estimate of voxel-wise signal and functional connectivity. The evaluated RTPs included slice-timing correction, motion correction, spatial smoothing, signal scaling, and regression of low-frequency fluctuations (high-pass filtering), motion parameters, motion derivatives, global signal, white matter/ventricle mean signals, and physiological noises modeled by the image-based retrospective correction of physiological motion effects (RETROICOR) [19] and the respiration volume per time (RVT) [20]. The examined noises included motion and physiological noises, which are the most prevalent noises in the fMRI signal and functional connectivity [21–25].

At evaluating the real-time processing performance, we should also consider the cost of computation time. If the benefit of an added process is negligible compared to its computational cost, there would be no need to use it in the limited time of rtfMRI. Also, since using more regressors in noise regression requires more scans before obtaining a usable signal to avoid overfitting [10], we should not include ineffective regressors in the real-time process. Thus, we evaluated the computation time of each processing and discussed the number of regressors as the cost of RTP.

In addition, we examined the advantages and disadvantages of different approaches of GLM for real-time analysis, such as cumulative GLM (cGLM) and incremental GLM (iGLM) [8]. The cGLM performs an ordinary regression at each volume using the whole history of available online data, while iGLM updates only the latest estimation without keeping data history. Most rtfMRI applications have used iGLM with its short computation time and small memory consumption. However, iGLM cannot update regressor values retrospectively. This limitation could be critical when a regressor is created online and can be improved by a retrospective update, such as high-pass filtering and physiological noise regressors. We demonstrated that this limitation of iGLM hinders real-time physiological noise regression performance so that cGLM is preferred in RTP when using physiological noise regression.

This study performed a series of simulation analyses structured as follows. Firstly, we compared the incremental and cumulative GLM regarding their quality of online-made regressors, such as high-pass filtering and physiological noise models. Since the result indicated that cGLM is preferred for RTP, the simulated RTP system implemented cGLM. We used this system for the real-time processing simulation with resting-state fMRI data and rtfMRI neurofeedback task data targeting the left amygdala activation (LA-NF) [26]. To evaluate RTP’s benefit in noise reduction, we calculated the amounts of variance explained by the motion and physiological noises in the real-time estimate. For the resting-state data, evaluation was conducted for voxel-wise signals and dynamic functional connectivity in the whole-brain regions. For the LA-NF task data, we conducted the noise analysis on the neurofeedback target signal and calculated the signals’ correlation between the RTP and offline process to measure the integrity of the neurofeedback signal.

## 2 Materials and Methods

### 2.1 Human Data

We performed the simulation analyses on fMRI resting-state data from 87 healthy participants and rtfMRI neurofeedback task data from 22 participants (14 major depressive disorder patients). The Supplementary material, ‘fMRI samples for the simulation analyses’ describes the details about the participants. All human neuroimaging data were acquired in the studies [27] and [26] approved by the Western Institutional Review Board, Puyallup, WA. Human research in this study was conducted according to the principles expressed in Declaration of Helsinki. All subjects gave written informed consent to participate in the study and received financial compensation.

### 2.2 Comparisons between the incremental and cumulative GLM

Most rtfMRI applications with regression use incremental GLM (iGLM) [8], which calculates the latest estimates (beta and residual) without keeping data history. Although this has a notable advantage in its short computation time and small memory requirements, iGLM cannot update past regressor values because the current estimation is made based on a previous estimate. This could be a disadvantage in real-time processing when the regressor values are formed online and the regressor model could be improved with increasing sample points, such as high-pass filtering and physiological noise regressors. The cumulative GLM (cGLM), in contrast, performs an ordinary regression at each sampling, using the whole history of available online data [6]. While the cGLM may take a longer computation time than iGLM, the online update of the past regressor values could improve the accuracy of the online-made regressors.

The aim of the present simulation analysis was to compare the quality of online-made regressors between the iGLM and cGLM approaches for high-pass filtering and physiological noise models. Specifically, we evaluated the frequency filtering performance of the high-pass filtering regressors and the accuracy of online-made physiological noise regressors of RETROICOR and RVT.

#### 2.2.1 High-pass filtering regression

The Legendre polynomial regressors and discrete cosine transform (DCT) basis sets have been used for high-pass filtering in AFNI (https://afni.nimh.nih.gov/) and SPM (https://www.fil.ion.ucl.ac.uk/spm/), respectively. We evaluated the frequency filtering performance of these regressors in iGLM and cGLM. We generated a random white-noise signal with a length of 250 samples, at a 2-s sampling interval and unit absolute power for all frequencies. GLM was applied to this signal with the Legendre polynomial or DCT regressors at 50, 100, 150, 200, and 250 lengths to simulate the real-time evaluation with a limited number of samples. Regressors in iGLM were made for the length of 250 TRs, and then the first part of them was extracted. This simulated that iGLM cannot update past regressor values—the same approach used in Bagarinao, Matsuo, Nakai and Sato [8]. In contrast, regressors in cGLM were updated at each length with the order adjustment (e.g., order of the Legendre polynomials or the cycles of DCT) for a designed pass frequency in each sampling length. The order of Legendre polynomials was calculated as 1 + floor(*d*/150), where *d* is the scan duration in seconds (default in AFNI). The threshold frequency of DCT was 1/128 Hz (default in SPM). The simulation was repeated 1,000 times with different random white noise signals. The frequency filtering performance was examined with the absolute power spectra of the residual signal.

We also performed the same simulation for resting-state fMRI signals to examine the difference of GLM approaches in frequency-filtering performance. The resting-state images of 87 participants (see Supplementary material ‘fMRI samples for the simulation analyses’) were applied slice-timing correction, motion alignment, and spatial smoothing using the RTP simulation system (Supplementary material ‘RTP simulation system implementation’). Then, the high-pass filtering regression was applied to the signal time-course of voxels within the intersection of the brain mask and a signal mask (3dAutomask in AFNI was applied to the functional images to make the signal mask). The signal in each voxel was normalized to z-score before applying the GLM. The average result of the whole-brain voxels for all participants is reported.

#### 2.2.2 Physiological noise model

RETROICOR and RVT regressors were evaluated in the online GLM comparison. We used AFNI’s RetroTS.py with the default parameters to make the regressors. RETROICOR regressors were Fourier basis sets with the phase synchronized to cardiac or respiration signal time-course [19]. A basis set of four regressors was made for each of the cardiac and respiration signals. RVT is a respiration volume per time calculated by detecting the top and bottom peaks of the respiration time-course to measure respiration volume [20]. The five time-shifted regressors were made.

The simulation examined how the online calculation of the RETROICOR and RVT regressors diverged from that in the offline calculation. We used respiration and cardiac signals recorded with resting-state fMRI (Supplementary material ‘fMRI samples for the simulation analyses’). The simulation was initiated at the time equal to 70 s in order to wait to obtain enough samples to create the regressors. In both GLM approaches, the regressors were recalculated at every TR (= 2 s) using the accumulated history of cardiac and respiration signals from the start of the scan until the current TR. There was no difference in the regressor calculation between the two GLM approaches. The only difference was the update method of regressor values at GLM; in the cumulative approach (cGLM), all values were replaced in the updated regressor values, while in the incremental approach (iGLM), only the latest TR value was updated, which is an inevitable limitation of iGLM. We used Pearson correlation between the online and offline regressors to evaluate the quality of the online regressor. Mean correlation across participants was calculated by applying Fisher’s z-transformation, averaging the transformed values, and then transforming back to the correlation coefficient.

### 2.3 Implementation of fMRI real-time processing simulation system

Figure 1 shows an overview of the simulation system. The WATCH module monitors a directory where a new fMRI volume file is created in real-time and loads it to send the data to the processing modules. In the simulation, we copied a pre-scanned image file volume-by-volume to the monitored directory. The RTP sequence included slice-timing correction with a temporal shifting of sampling time (TSHIFT), motion correction with volume registration (VOLREG), spatial smoothing by convolving Gaussian kernel within a mask (SMOOTH), and noise regression (REGRESS) with cGLM. Signal scaling is done within the REGRESS module. Details of each process are in the Supplementary material (RTP simulation system implementation). The code for the simulation system is available on GitHub (https://github.com/mamisaki/fMRI_RTP_Simulation).

**Figure 1.**
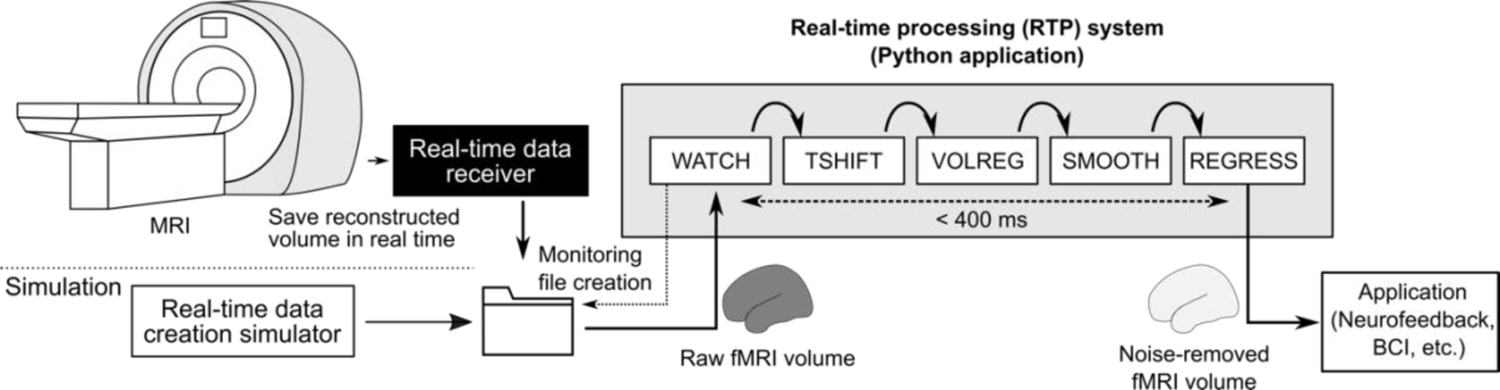
Overview of the fMRI real-time processing simulation system. The system monitors a directory where an fMRI data volume file is created in real-time. The WATCH module detects new file creation and reads the data to send it to a process sequence of TSHIFT (slice-timing correction), VOLREG (motion alignment), SMOOTH (spatial smoothing) and REGRESS (scaling and general linear model analysis). The output (residual of the noise regression as a noise-removed fMRI signal) can be sent to an independent application of neurofeedback or a brain-computer interface. The simulation was performed by copying a volume image of a pre-scanned fMRI data into a monitored directory.

### 2.4 Real-time processing simulation and offline analysis

The simulation was performed on a Linux computer (Ubuntu 16.04 LTS) with dual Intel Xeon CPUs (Gold 6126, 2.6 GHz, 12 cores for each), 256 GB RAM, and an NVIDIA TITAN V GPU (5120 CUDA cores with 12 GB memory). The simulation was started by copying an fMRI volume file in the directory monitored by the WATCH module. The resting-state fMRI data of 87 participants and the rtfMRI neurofeedback task data of 22 participants were used in the simulation. Their details are described in the Supplementary material, ‘fMRI samples for the simulation analyses.’

We tested ten types of RTP pipelines to evaluate the benefit of each processing step and regressor. Table 1 shows the process and regressors included in each pipeline. RTP0 included only the volume registration (VOLREG) for motion correction, a conventional minimum RTP, used as a baseline of the evaluation. RTP1 added slice-timing correction (TSHIFT) before VOLREG, and RTP2 added spatial smoothing (SMOOTH) to RTP1. From RTP3 to RTP9, noise models were regressed out. The regressors added at each RTP were high-pass filtering (HPF) with Legendre polynomials in RTP3, motion parameters (Mot) of three shifts and three rotations in RTP4, temporal derivatives of the motion parameters (dMot) in RTP5, global signal (GS, mean signal within a brain mask) in RTP6, white matter/ventricle mean signals (WM, Vent) in RTP7, RETROICOR (four cardiac and four respiration basis sets) in RTP8, and RVT (respiration volume per time with five time-shifted regressors) in RTP9. Details of each process and regressor are described in the Supplementary material (RTP simulation system implementation). The regression was done with cGLM. Residual volumes of the regression were used as the processed signal [17]. We also ran offline processing using AFNI (details are shown in the Supplementary material, ‘Offline fMRI processing’).

**Table 1.** Real-time processing (RTP) pipelines.

| RTP pipeline | TSHIFT | VOLREG | SMOOTH | REGRESS |  |  |  |  |  |  |
| --- | --- | --- | --- | --- | --- | --- | --- | --- | --- | --- |
|  |  |  |  | High-pass filtering | Motion | Motion derivative | Global signal | WM/Vent means | RICOR | RVT |
| RTP0: VOLREG |  | ✓ |  |  |  |  |  |  |  |  |
| RTP1: +TSHIFT | ✓ | ✓ |  |  |  |  |  |  |  |  |
| RTP2: +SMOOTH | ✓ | ✓ | ✓ |  |  |  |  |  |  |  |
| RTP3: +REG[HPF] | ✓ | ✓ | ✓ | ✓ |  |  |  |  |  |  |
| RTP4: +REG[Mot] | ✓ | ✓ | ✓ | ✓ | ✓ |  |  |  |  |  |
| RTP5: +REG[dMot] | ✓ | ✓ | ✓ | ✓ | ✓ | ✓ |  |  |  |  |
| RTP6: +REG[GS] | ✓ | ✓ | ✓ | ✓ | ✓ | ✓ | ✓ |  |  |  |
| RTP7: +REG[WM, Vent] | ✓ | ✓ | ✓ | ✓ | ✓ | ✓ | ✓ | ✓ |  |  |
| RTP8: +REG[RETROICOR] | ✓ | ✓ | ✓ | ✓ | ✓ | ✓ | ✓ | ✓ | ✓ |  |
| RTP9: +REG[RVT] | ✓ | ✓ | ✓ | ✓ | ✓ | ✓ | ✓ | ✓ | ✓ | ✓ |

VOLREG, volume registration for motion correction; TSHIFT, slice-timing correction; SMOOTH, spatial smoothing with convolving Gaussian kernel (FWHM=6mm); REGRESS (REG), regressing out a noise time-course; WM, white matter; Vent, ventricle; RVT, respiration volume per time.

The computation time of each RTP was measured by the time from receiving a volume from the previous step to sending the processed volume to the next step in the simulation system. The start time of the WATCH module was the file creation time.

### 2.5 Noise amount in the real-time processed signals

#### 2.5.1 Evaluated signals

For the resting-state data, we evaluated the noise amount in an RTP voxel-wise signal and dynamic functional connectivity (FC) time-course. A linear trend was removed from the voxel-wise signals of RTP0, RTP1, and RTP2. Dynamic FC was calculated for the detrended signal. For the RTP3 and the later pipelines, detrending was included in the high-pass filtering regression. Dynamic FC was calculated for all combinations of the 264 functional ROIs (6mm-radius spheres) defined by Power, Cohen, Nelson, Wig, Barnes, Church, Vogel, Laumann, Miezin, Schlaggar and Petersen [28]. The ROIs in the MNI template space were warped into the participant’s native image space using the Advanced Normalization Tools (ANTs, http://stnava.github.io/ANTs/).

We used two methods to calculate the dynamic FC; sliding-window correlation [29] and two-point algorithm [30]. The sliding-window correlation is a z-transformed Pearson correlation between ROIs within a time window [31]. A rectangular window of 5-TR width was used in the simulation. The window was moved at each TR to calculate the time-course of dynamic FC. The two-point algorithm assesses the agreement of change directions between ROIs. The feedback signal was given a binary value with positive reinforcement feedback (+1) when the two regions had the same change direction and no reinforcement feedback (0) when the change directions were different (if we wanted to train a participant to increase the connectivity). While the original introduction of the two-point method [30] used another ROI as a control to cancel a signal change unspecific to the target connectivity, we did not use a control ROI in the current simulation because another study [32] showed a control ROI is unnecessary when comprehensive noise reduction is applied.

For the neurofeedback task data, the mean signal in the left amygdala region, which was the target signal in the self-regulation task [26], was extracted for the noise evaluation. We applied the same procedures as for the voxel-wise signals of the resting-state data. The left amygdala mask was defined in the MNI template brain with FreeSurfer (https://surfer.nmr.mgh.harvard.edu/) segmentation and warped into an individual brain space using ANTs.

#### 2.5.2 Noise variance ratio

The motion and physiological noise amounts in the RTP signal were measured by a variance ratio explained by motion, cardiac, and respiration noise time-courses. Specifically, we performed a multiple regression analysis for the RTP voxel-wise and dynamic FC signals with noise regressors to calculate the coefficient of determination. The coefficient of determination with noise regressors, *R*_n_^2^, indicates the variance ratio explained by noises. We used this measure as the noise amount in the RTP signal. The first 65 TRs (62 TRs plus three TRs before the steady state) were excluded from the regression analysis because the output of real-time regression with a small number of samples is unreliable due to overfitting [10].

The effect of the motion, cardiac, and respiration noises were evaluated independently. Noise regressors were calculated in an offline process. The motion noise regressor was a time-course of frame-wise displacement, calculated by the root sum of squared motion temporal derivatives. The cardiac noise regressors included heart rate (HR) [33] and the RETROICOR model [19]. The HR time-course was calculated using BioSPPy library in python (https://biosppy.readthedocs.io/en/stable/) from a pulse oximetry measurement, resampled at each TR, and convolved with the cardiac response function presented in Chang, Cunningham and Glover [33]. The RETROICOR regressors were calculated offline using AFNI’s RetroTS.py. The four basis regressors for the cardiac signal were extracted.

The respiration noise regressors included the respiration variance (RV) [33, 34], the RETROICOR model, and RVT. The RV is a variance of respiration signal within a 3s-window centered at each TR [33]. The RV time-course was convolved with the respiration response function shown in Chang, Cunningham and Glover [33] and provided by Power, Lynch, Dubin, Silver, Martin and Jones [34]. The RETROICOR and RVT regressors were calculated offline using AFNI’s RetroTS.py. The four basis regressors for the respiration signal and the five RVT regressors were used.

In the noise regression for dynamic FC, the windowed calculation could obscure the temporal association between the noise regressors and the dynamic FC. For example, in the sliding-window dynamic FC, any motions within the time window could affect the current neurofeedback signal. However, if we evaluated only the immediate association between the noise and RTP signal, we could miss a delayed effect of the motion within a window. Therefore, we added windowed regressors, the mean and standard deviation of the noise regressors within the connectivity calculation window, to the noise regressors. For example, the motion noise regressors for the dynamic FC were composed of the frame-wise displacement (FD) at each TR and the sliding-window mean and standard deviation of the FD within the FC calculation window (five TRs for the sliding-window and two TRs for the two-point method). If the node ROI signals were both affected by the same noise, their dynamic FC could increase when the standard deviation of the noise within the window was high. When the noise regressor is a variance measure (e.g., respiration variance), dynamic FC could associate with the mean noise regressor value within the window. We included these possible delayed noise effects in the present noise analysis.

We also investigated the group-level association for participant-wise mean connectivity with the motion and physiological noises. Weiss, Zamoscik, Schmidt, Halli, Kirsch and Gerchen [18] indicated that subject-wise mean FC feedback signal could be correlated with breath rate. To investigate how RTP could eliminate this confounding effect, we calculated the mean dynamic FC across time and ROIs, and examined its association with each participant’s mean framewise displacement, standard deviation of heart rate, and standard deviation of respiration rate. Heart rate and respiration rate were calculated for each TR using the BioSPPy library, and then their standard deviations in a scan run were calculated for each participant.

### 2.6 Integrity of the neurofeedback signal

For the neurofeedback task data, we evaluated the benefit of RTP in improving the integrity of the neurofeedback signal. Integrity, in this case, refers to how well the signal reflects neural activation, free from noises. Although the signal amplitude relative to a rest block or the temporal contrast to noise ratio (TCNR) could be considered a measure of signal quality, we argue that these do not necessarily indicate the RTP benefit. For example, if the motion or physiological noise correlated with a self-regulation task, the task-related signal change could be larger for an RTP without noise corrections. Indeed, such a task-noise association has been reported in Weiss, Zamoscik, Schmidt, Halli, Kirsch and Gerchen [18]. TCNR could also be higher for an RTP with fewer noise corrections if a noise changed with the task time course. Such noise could reduce random signal fluctuation and increase the TCNR.

Since the size of task-related signal change cannot be a reliable measure of neurofeedback signal integrity, we used the signal correlation between RTP and offline processing to evaluate the RTP benefit. Assuming that the offline processed signal has the best available quality, the RTP signal with a higher correlation with the offline processed one should have better neurofeedback signal integrity. We calculated the Pearson correlation between the RTP and offline processed signals in the left amygdala for each participant, applied Fisher’s z-transformation and averaged across the participants for each RTP pipeline.

### 2.7 Statistical analysis

Participant-wise average *R*_n_^2^ across the brain voxels or connectivity was compared between the RTP pipelines. Linear mixed-effect model analysis was applied to the average^2^ with a fixed effect of the pipeline and a random effect of the participant on the intercept. We used the *lme4* [35] with the *lmerTest* package [36] in R language and statistical computing [37]. The ad-hoc comparison between pipelines was performed in a sequential way to test the individual effect of the real-time process and regressor (e.g., RTP1-RTP0 for evaluating the TSHIFT effect, RTP2-RTP1 for evaluating the SMOOTH effect, RTP3-RTP2 for evaluating the high-pass filtering regressor effect). The comparison was performed with the *lsmeans* package [38] on R with multiple testing correction through the multivariate *t*-distribution method.

We also tested the noise effect in each voxel and connectivity. Since the BOLD signal, as well as the physiological signals, includes an autocorrelation component, a statistical test with a null hypothesis assuming the Gaussian noise could underestimate the false positive rate [39]. Thus, we used the phase randomization test [39, 40] to evaluate the statistical significance of the *R*_n_^2^ value. Phase-randomization was applied to the noise regressors. For the dynamic FC, the windowed average and standard deviation regressors were calculated after the phase-randomization. The randomized regressors were made 5,000 times for each participant.

All the noise calculation was done in the participant’s native space because resampling in spatial normalization could affect the signal time series and change the noise contribution [23]. For the group analysis of the voxel-wise signal, the individual participant’s^2^ values were warped into the MNI template brain to calculate the mean *R*_n_ ^2^ across participants in each voxel. *R*_n_^2^ maps of phase-randomized regressors were also warped into the MNI to make a null distribution of the mean *R*_n_^2^ for calculating *p*-value. The false discovery rate (FDR) correction with the Benjamini-Hochberg procedure was applied to the *p*-values. The maps were thresholded with FDR < 0.05.

For the neurofeedback task data, we used a permutation test to evaluate the statistical significance of the *R*_n_ ^2^ value for each pipeline and comparisons between the pipelines, and FDR correction was applied to *p*-values. We used the same linear mixed-effect model as for the resting-state data to test the correlation between RTP and offline processed signals.

## 3 Results

### 3.1 Comparisons between the incremental and cumulative GLM

Figure 2 shows the high-pass filtering performance of Legendre polynomial and DCT regressors for the white noise signal (Fig. 2A) and the resting-state fMRI signal (Fig. 2B). Figure 2A indicates that frequencies higher than the designed threshold (vertical dotted line) decreased more with iGLM than with cGLM, especially when the number of TRs was short, demonstrating that iGLM had a less accurate high-pass filtering property. Figure 2B shows a similar tendency for the BOLD signal, while the difference between the GLM approaches was less prominent than for white noise.

**Figure 2.**
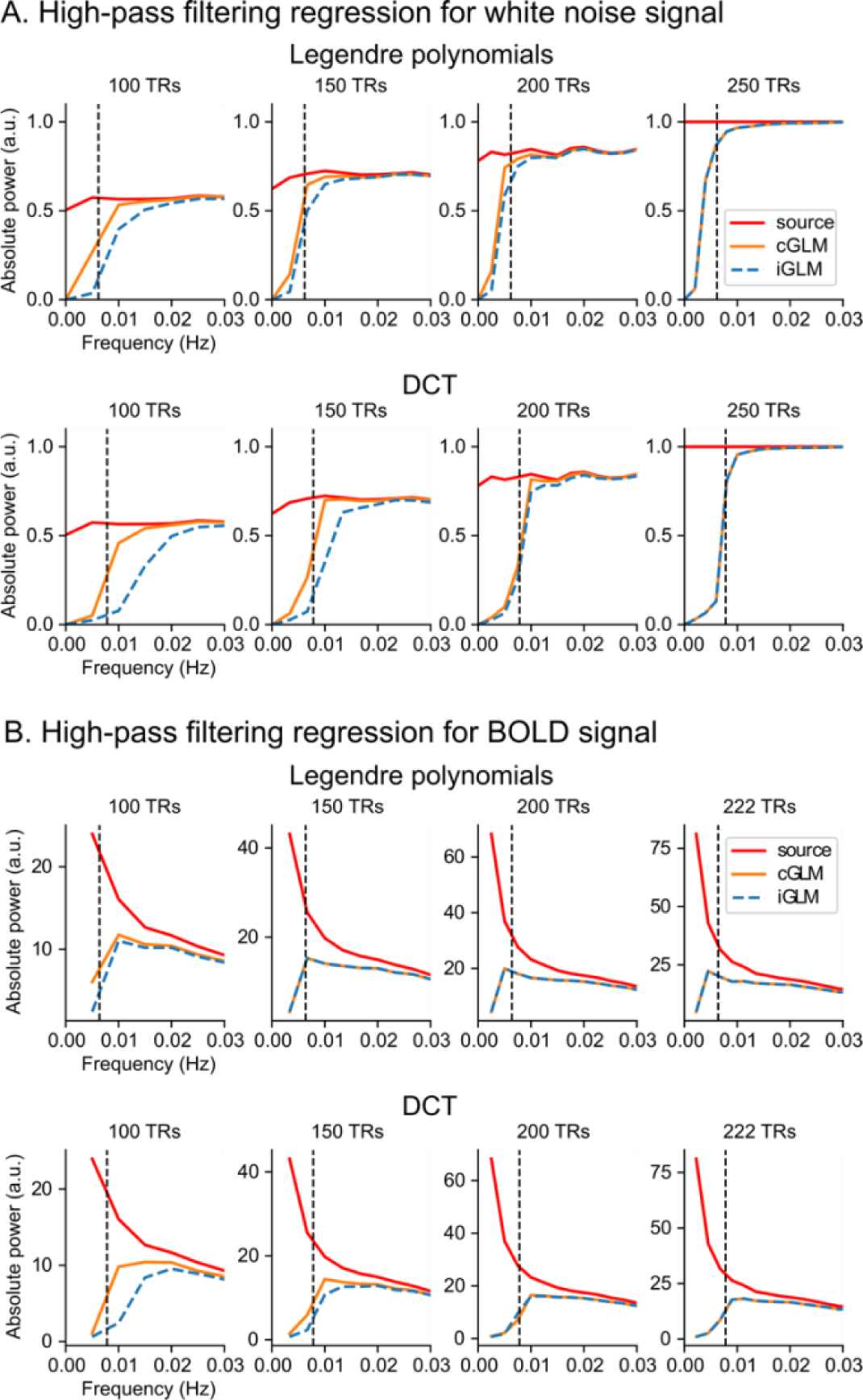
Comparisons between the incremental and cumulative GLM in high-pass filtering. The absolute frequency powers of filtered signal with incremental GLM (iGLM) and cumulative GLM (cGLM) are shown. The red line is the power spectrum of the source signal and the orange and blue lines are power spectrums of the filtered signals with high-pass filtering regression using iGLM and cGLM, respectively. The vertical dotted line is the designed pass-frequency threshold. A. Random white noise signal was regressed with Legendre polynomial regressors and discrete cosine transformation basis sets (DCT). The plots are the average of 1,000 times of simulation. B. BOLD signals in resting-state fMRI were regressed with Legendre polynomial regressors and discrete cosine transformation basis sets (DCT). The plots are the average of the whole brain voxels for 87 participants.

Figure 3 shows the correlation between the online and offline calculations of the RETROICOR and RVT regressors for the incremental (iGLM) and cumulative (cGLM) approaches. No difference between iGLM and cGLM was observed for the cardiac regressors (Card). For the respiration regressors (Resp), cGLM had a higher correlation with the offline calculation. For the RVT regressors, iGLM had considerably lower correlations with the offline calculation. This indicates that online-made physiological noise regressors were inaccurate in iGLM without retrospective correction, especially for the respiration and RVT regressors. These results suggest that cGLM is preferred to iGLM in real-time processing, as far as computation time permits.

**Figure 3.**
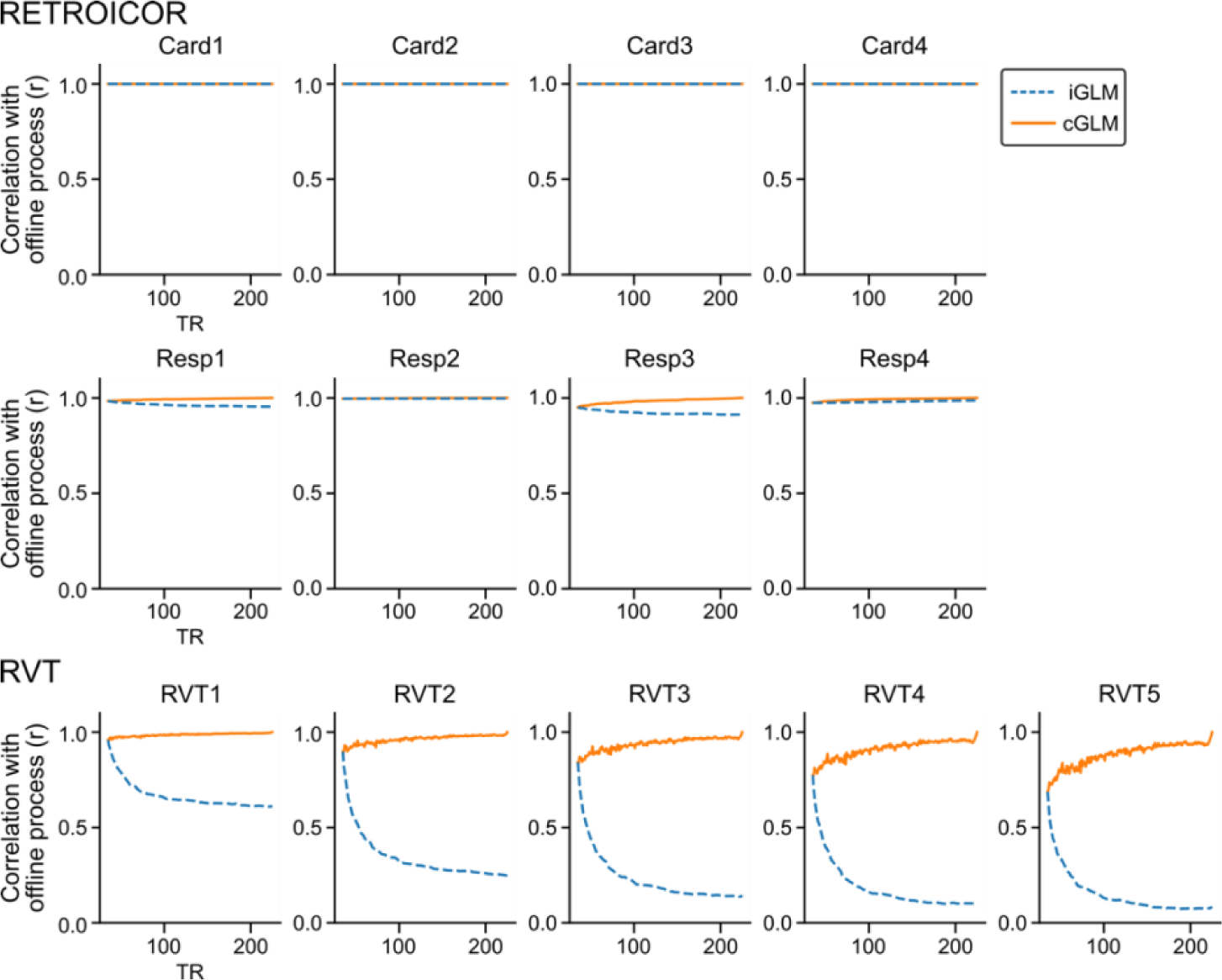
Comparisons between the incremental and cumulative GLM’s online-made physiological noise regressors for their correlation with the offline-made regressors. Correlations are shown for RETROICOR (respiration [Resp] and cardiac [Card] basis sets) and RVT regressors. The lines are the average for 87 participants.

### 3.2 Computation time

Figure 4 shows the average computation time across participants for each processing module at each TR. The processing was applied to whole-brain voxels (149082 voxels on average). The REGRESS process used the cumulative approach and included all the regressors implemented in the system. The REGRESS process started at the 42nd TR to allow for the acquisition of enough samples for the regression (the initial three volumes were excluded from the process to ensure the fMRI signal reached the steady state; thus, 39 samples were available at the 42nd TR).

**Figure 4.**
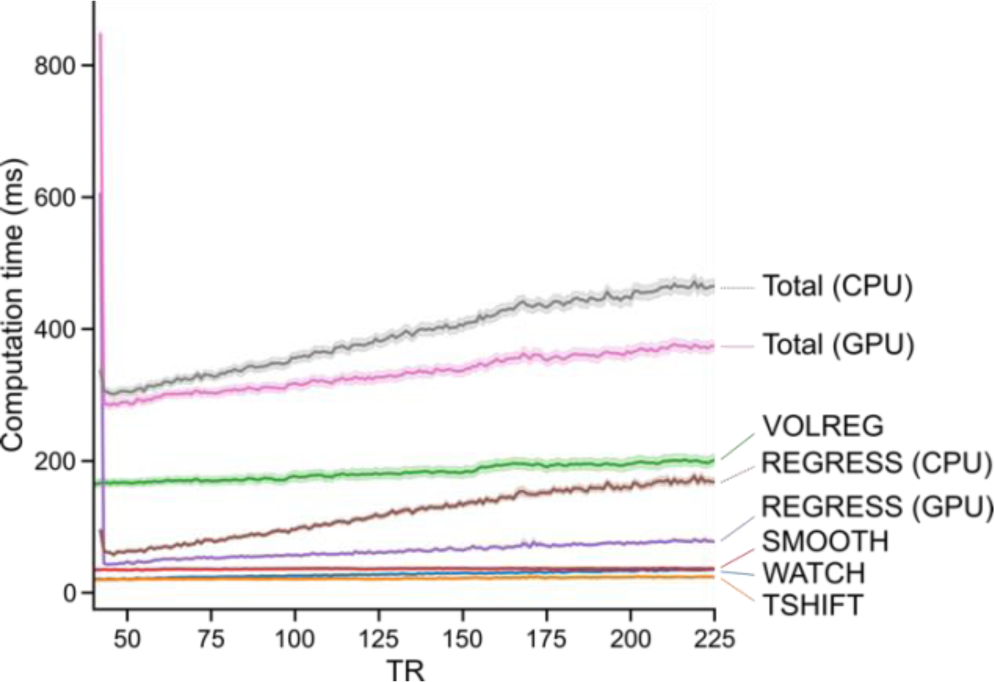
Computation times of each RTP module. The mean computation time across participants at each TR is shown for each processing module with a 95% confidence interval (band around the line). The time was between receiving a volume from the previous step and sending the processed volume to the next step. The start time of the WATCH module was the file creation time in the monitored directory. REGRESS performed cumulative GLM and included all the regressors implemented in the system (Legendre polynomials, motion parameters, motion derivatives, global signal, white matter/ventricle mean signals, RETROICOR, and RVT).

The REGRESS’s initial processing TR took a long time due to the initialization process (see Supplementary material, ‘RTP simulation system implementation’). Other than the initial TR of REGRESS, the most time-consuming processing was the VOLREG, volume registration for motion correction. We found resampling the volume image in the aligned grid took a long time in VOLREG. The time for REGRESS with CPU increased with TR, which was because cumulative calculation used more data in a later TR. The slope of the increase was less steep with GPU computation. Other processes took less than 100 ms each, and the total process time was less than 400 ms with GPU and less than 500 ms with CPU, which would be short enough for a majority of rtfMRI applications.

### 3.3 Noise evaluation in the real-time processed signal for resting-state data

#### 3.3.1 RTP effects on the noise variance ratio (*R*_n_^2^) in the voxel-wise signal

Figure 5A shows distributions of the mean noise variance ratio (*R*_n_^2^) explained by the motion and physiological noises for the voxel-wise signal. The plot shows the distribution of the participant mean values in the brain. The voxels in the white matter and the ventricle areas were excluded from the mean calculation (the masks for the white matter/ventricles signal regression in RTP were used). Significant decreases or increases of the mean noise variance ratio by adding an RTP component were summarized in Table 2 (statistical values for each comparison are shown in Supplementary Table S1).

**Figure 5.**
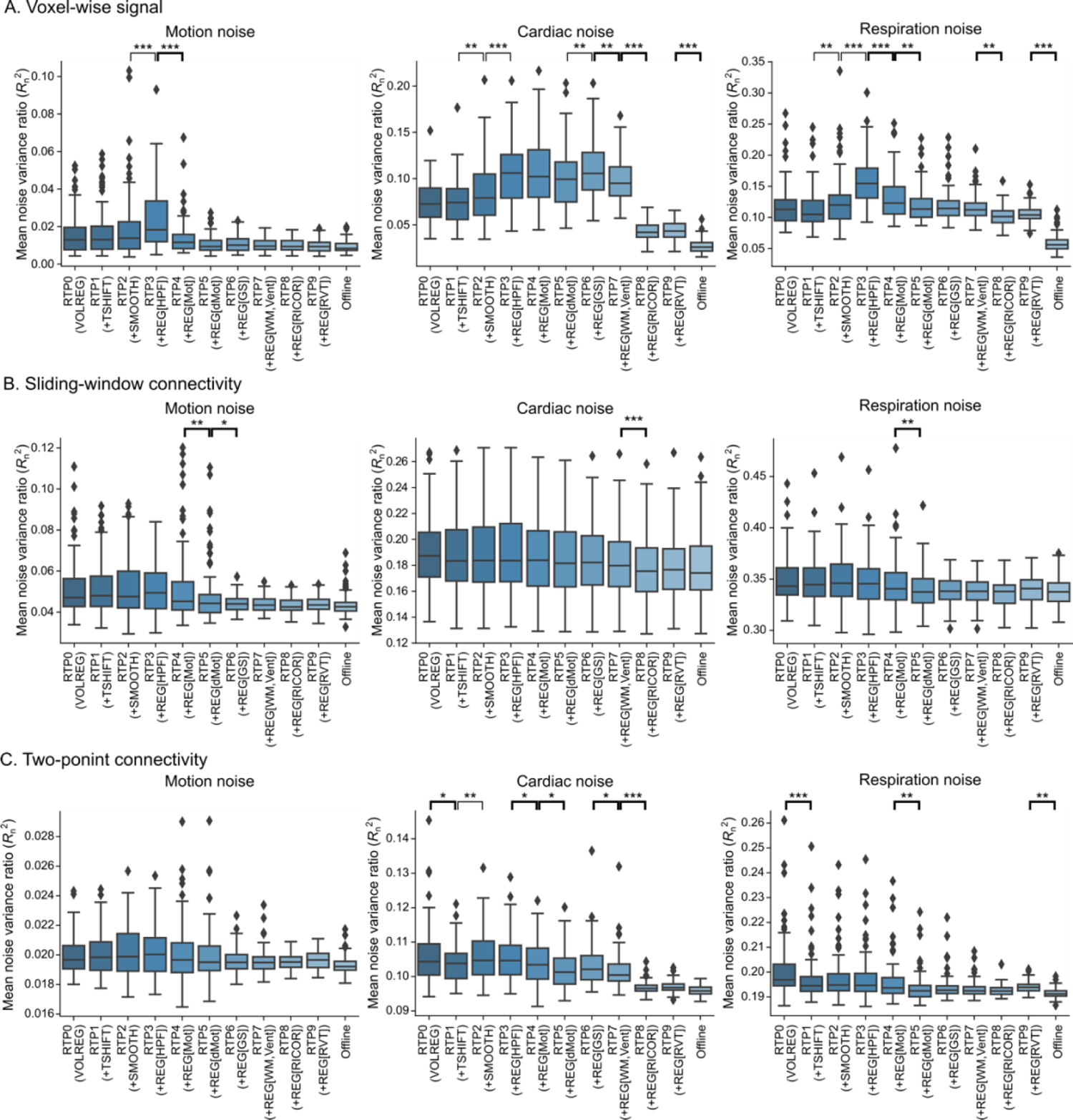
Box plot of the mean *R_n_*^2^ of noise regression on voxel-wise signals (A), sliding-window (5-TR width) connectivity (B), and two-point connectivity (C). The lines with stars indicate the significant difference in the mean *R_n_*^2^ between the pipelines; thin lines indicate increased and thick lines indicate decreased noise variance ratio. HPF, high-pass filtering regressor; Mot, motion regressor; dMot, motion derivative regressor; GS, global signal regressor; WM, Vent, white matter/ventricle mean signals regressors; RICOR, RETROIOCR regressor; RVT, respiration volume time-course regressor; *, *p* < 0.05; **, *p* < 0.01; ***, *p* < 0.001.

**Table 2.** Summary of the RTP noise reduction performance for voxel-wise signals

| RTP | Noise |  |  |
| --- | --- | --- | --- |
|  | Motion | Cardiac | Respiration |
| VOLREG+TSHIFT | - | - | - |
| +SMOOTH | - | $\Delta^{**}$ | $\Delta^{**}$ |
| +REG[HPF] | $\bigcirc^{***}$ | $\Delta^{***}$ | $\bigcirc^{***}$ |
| +REG[Mot] | $\bigcirc^{***}$ | - | $\bigcirc^{***}$ |
| +REG[dMot] | - | - | - |
| +REG[GS] | - | $\Delta^{**}$ | - |
| +REG[WM, Vent] | - | $\bigcirc^{**}$ | - |
| +REG[RETROICOR] | - | $\bigcirc^{***}$ | $\bigcirc^{**}$ |
| +REG[RVT] | - | - | - |

The cardiac and respiration noise ratios increased with spatial smoothing (SMOOTH, △ in Table 2). While high-pass filtering regression (REG[HPF]) also increased the cardiac noise ratio, it significantly reduced the motion and respiration noise ratios (ϒ in Table 2). Further reduction of the mean noise variance ratio was seen with the regression of motion parameters (REG[Mot]) for the motion and respiration noises, the regression of mean white matter/ventricle signals (REG[WM, Vent]) for the cardiac noise, and RETROICOR (REG[RICOR]) for the cardiac and respiration noises. Global signal regression (REG[GS]) increased the mean cardiac noise ratio. Interestingly, no reduction of respiration noise was observed with the RVT regressor (REG[RVT]) despite the RVT model included in the respiration noise regression analysis.

Voxel-wise analysis results are shown in figures 6, 7, and 8. Figure 6 shows maps of the voxel-wise significant motion noise *R*_n_^2^. Adding the motion parameter regression removed most of the significant voxels (RTP4), and adding the motion derivative regression (RTP5) eliminated a significant motion noise ratio in all voxels. Supplementary Figure S1 shows significant differences in the motion noise *R*_n_^2^ at each sequential contrast. TSHIFT and SMOOTH increased the motion noise ratio in many voxels, especially around the white matter and the ventricle areas. A significant decrease was seen when adding the regressions of high-pass filtering (REG[HPF]), motion parameters (REG[Mot]), motion derivatives (REG[dMot]), and the mean white matter/ventricle signals (REG[WM, Vent]).

**Figure 6.**
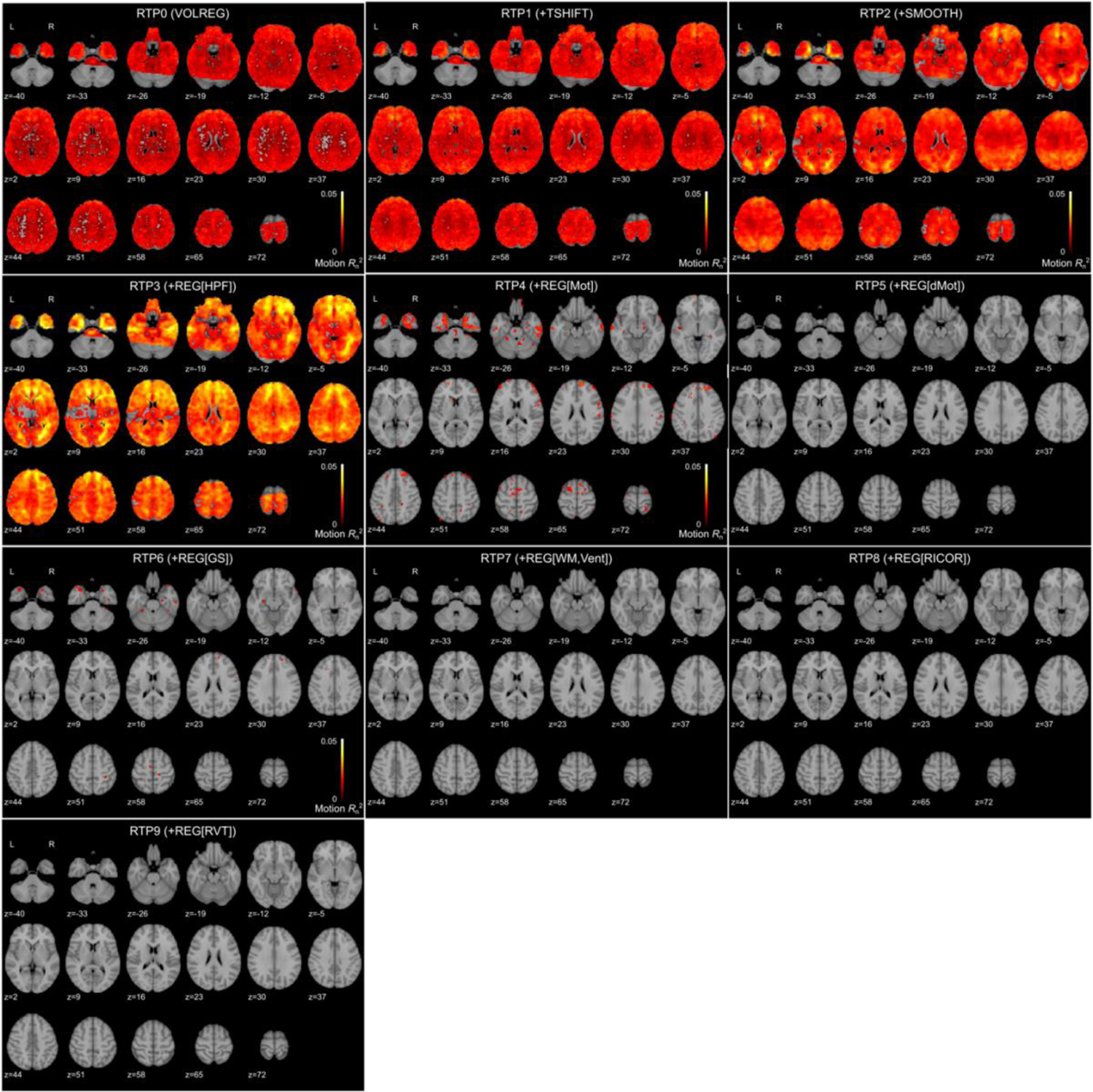
Significant *R_n_* ^2^ of motion noise in the voxel-wise signal for each real-time processing pipeline. The maps were thresholded by FDR < 0.05 with a randomization test.

Figure 7 shows maps of the voxel-wise significant cardiac noise *R*_n_^2^. Without RETROICOR, a significant cardiac noise ratio was seen in the whole-brain area and was especially high in the medial to anterior temporal and the anterior cingulate areas, overlapped with the large cerebral blood vessels. Adding the RETROICOR regression (RTP8) removed most of the significant voxels, although a significant effect remained in the large blood vessel areas. Supplementary Figure S2 shows significant differences in the cardiac noise *R*_n_^2^ (FDR < 0.05) at each sequential contrast. Spatial smoothing (SMOOTH), high-pass filtering regression (REG[HPF]), and global signal regression (REG[GS]) significantly increased the cardiac noise ratio in many voxels. In contrast, significant decreases were seen by adding the regressors of motion derivatives (REG[dMot]), white matter/ventricle signals (REG[WM, Vent]), and the RETROICOR (REG[RICOR]). The RETROICOR regression had the most substantial effect among them.

**Figure 7.**
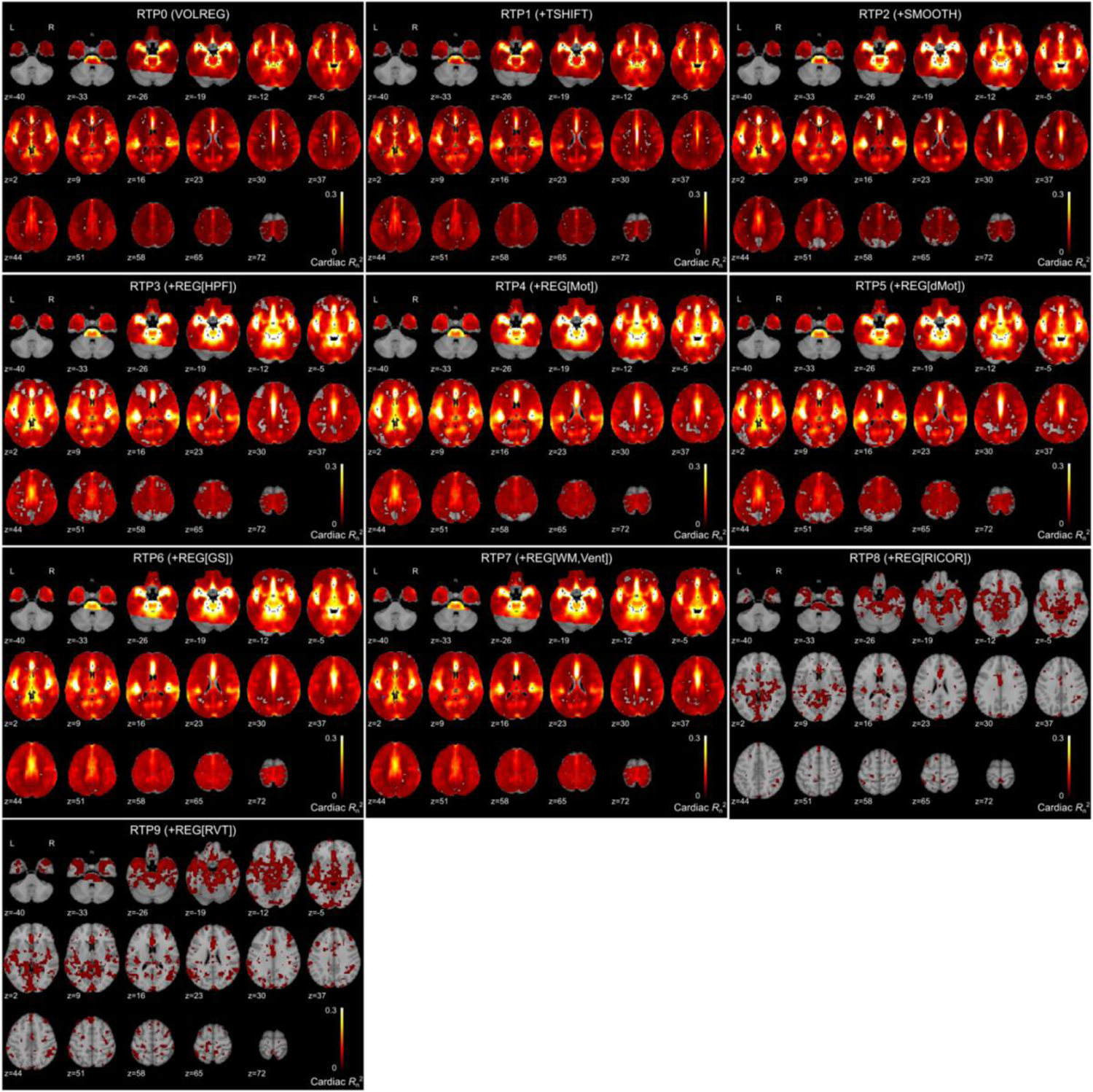
Significant *R_n_* ^2^ of cardiac noise in the voxel-wise signal for each real-time processing pipeline. The maps were thresholded by FDR < 0.05 with a randomization test.

Figure 8 shows maps of the voxel-wise significant respiration noise *R*_n_^2^. Without RETROICOR, a significant respiration noise ratio was seen in the whole-brain region, and most of the significant voxels disappeared by adding RETROICOR. Supplementary Figure S3 shows significant differences of the respiration noise *R*_n_^2^ at each sequential contrast. Spatial smoothing (SMOOTH) and high-pass filtering regression (REG[HPF]) increased the respiration noise ratio in the white matter and around the ventricle areas. A significant decrease was seen when adding the regressors of motion parameters (REG[Mot]), motion derivatives (REG[dMot]), white matter/ventricle signals (REG[WM, Vent]), and the RETROICOR (REG[RICOR]). Interestingly, adding the RVT regression increased the respiration noise ratio significantly in many voxels.

**Figure 8.**
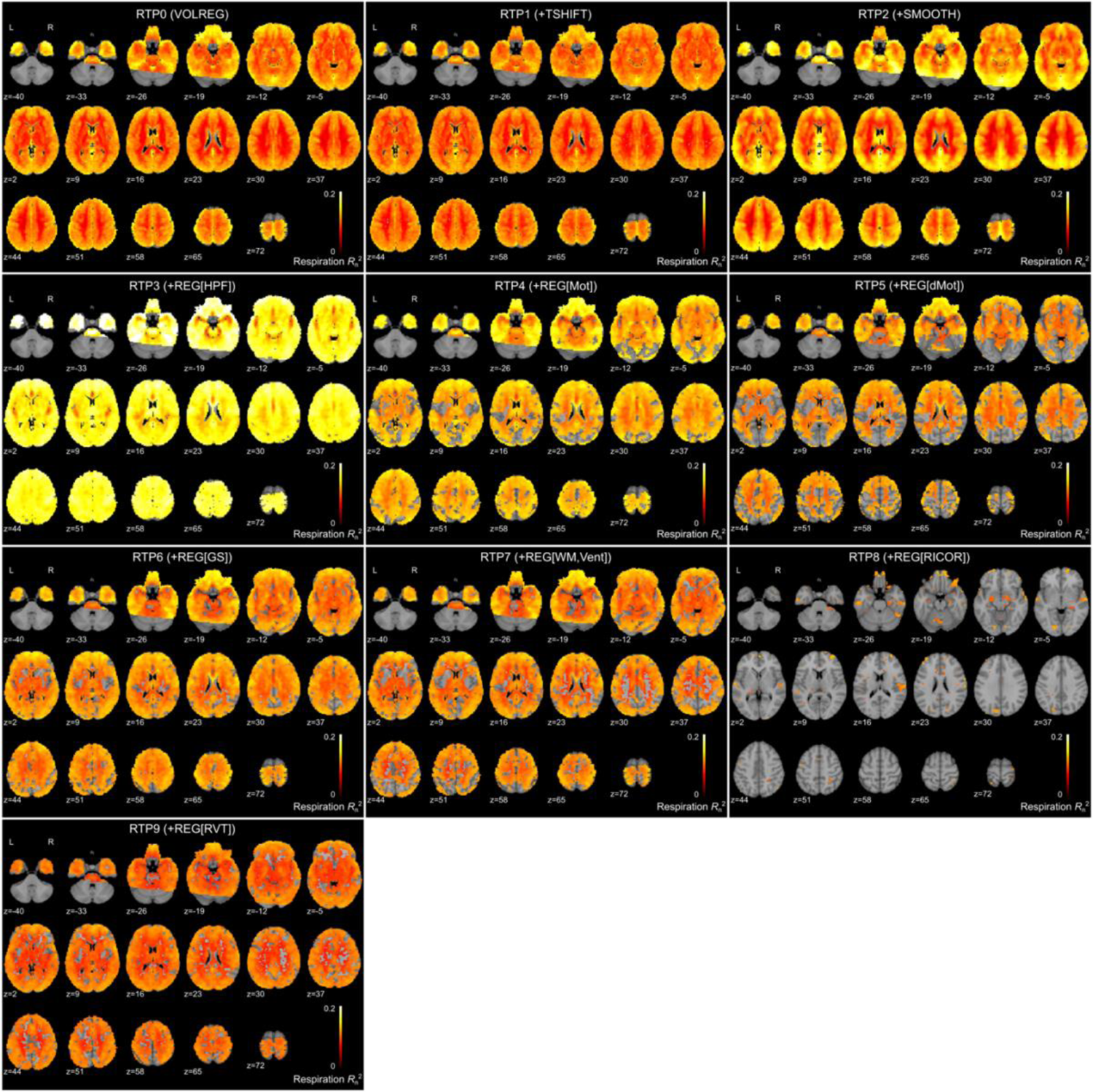
Significant *R_n_* ^2^ of respiration noise in the voxel-wise signal for each real-time processing pipeline. The maps were thresholded by FDR < 0.05 with a randomization test.

#### 3.3.2 RTP effects on the noise variance ratio (*R*_n_^2^) in the sliding-window connectivity

Figure 5B shows distributions of the mean noise variance ratio (*R*_n_^2^) explained by the motion and physiological noises for the 5-TR sliding-window dynamic FC time-series. The plot shows the distribution of participant mean values of all pairwise connectivity between the 264 functional ROIs. A significant decrease in the noise ratio by adding an RTP component is summarized in Table 3 (statistical values for each comparison are shown in Supplementary Table S2). The motion derivative regression (REG[dMot]) significantly reduced the mean motion and respiration noise ratios. Global signal regression (REG[GS]) reduced the mean motion noise ratio, and RETROICOR (REG[RICOR]) reduced the mean cardiac noise ratio.

**Table 3.** Summary of the RTP noise reduction performance for the sliding-window connectivity time-<u>course</u>

| RTP | Noise |  |  |
| --- | --- | --- | --- |
|  | Motion | Cardiac | Respiration |
| VOLREG+TSHIFT | - | - | - |
| +SMOOTH | - | - | - |
| +REG[HPF] | - | - | - |
| +REG[Mot] | - | - | - |
| +REG[dMot] | ○** | - | ○** |
| +REG[GS] | ○* | - | - |
| +REG[WM, Vent] | - | - | - |
| +REG[RETROICOR] | - | ○** | - |
| +REG[RVT] | - | - | - |

Connectivity-wise analysis results are shown in figures 9 and 10. Figure 9 shows maps of the connectivity-wise significant motion noise *R*_n_^2^. A significant motion noise was seen across whole-brain connectivity before adding the motion derivative regressor (REG[dMot]). After adding REG[dMot] (RTP5), no significant motion noise ratio was seen in any connectivity. No connectivity-wise significant difference of the motion noise *R*_n_^2^ was seen in any connectivity at any sequential contrast.

**Figure 9.**
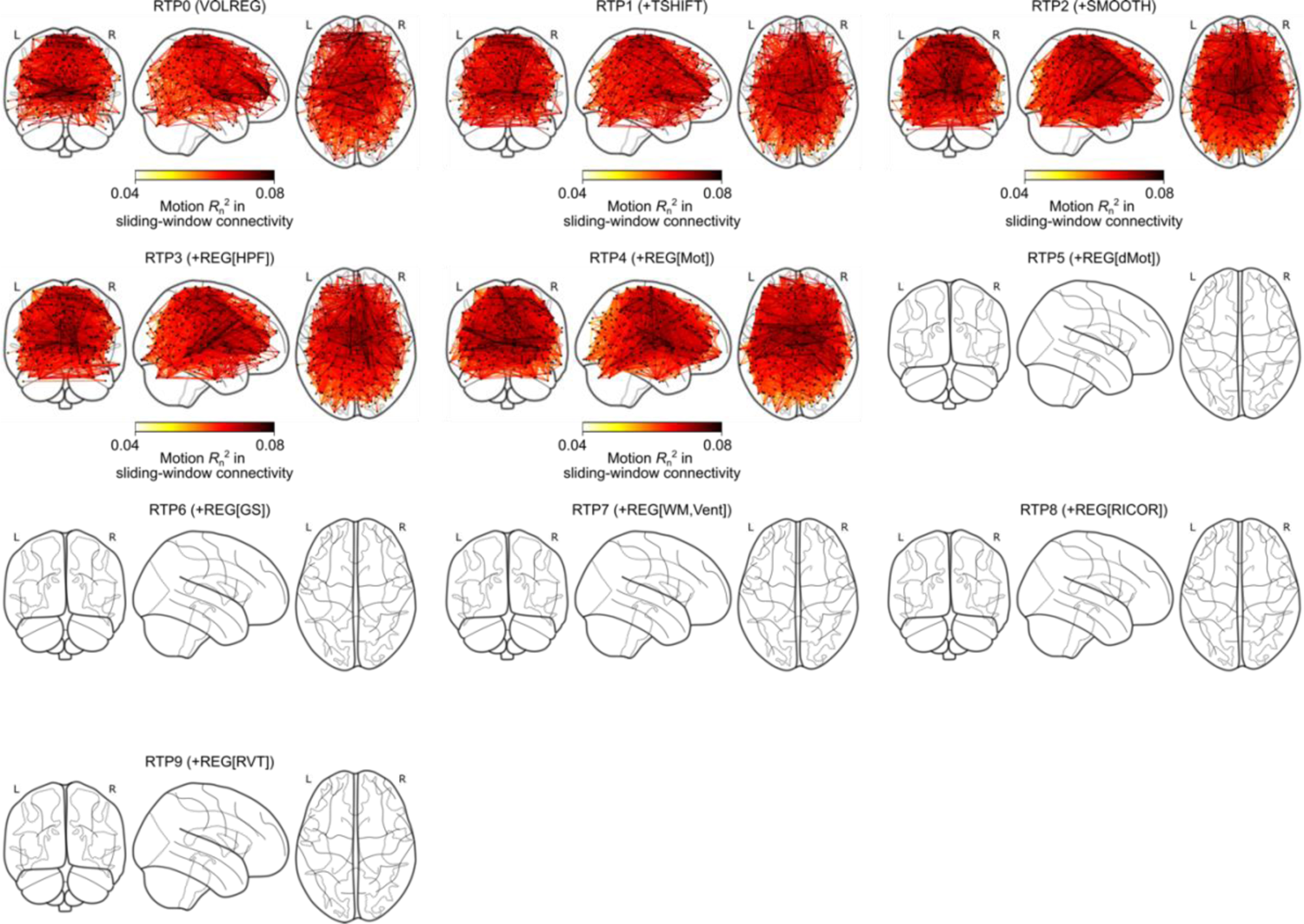
Connectivity plots of significant *R_n_*^2^ of motion noise in sliding-window (5-TR width) connectivity time-course for each real-time processing pipeline. The plots were thresholded by FDR < 0.05 with randomization test.

**Figure 10.**
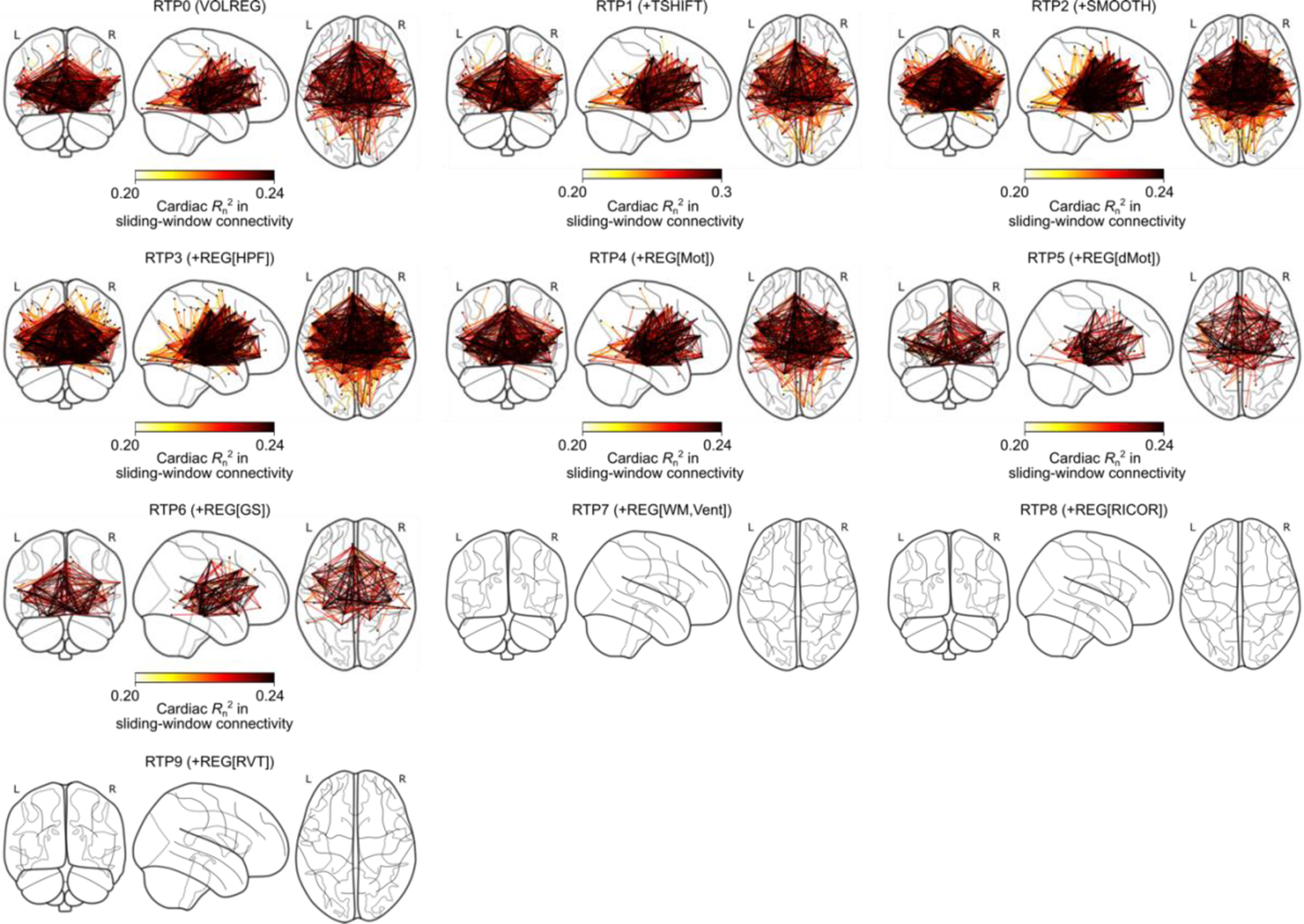
Connectivity plots of significant *R_n_*^2^ of cardiac noise in sliding-window (5TR-width) connectivity time-course for each real-time processing pipeline. The plots were thresholded by FDR < 0.05 with a randomization test.

Figure 10 shows maps of the connectivity-wise significant cardiac noise *R*_n_^2^. A significant cardiac noise was seen in the connectivity across the anterior to medial temporal and the anterior cingulate areas. Connectivity in the inferior occipital area also showed a significant cardiac noise ratio. The significant cardiac noise ratio gradually decreased with adding the regressors of motion parameters (REG[Mot]), motion derivatives (REG[dMot]), and global signal (REG[GS]). After adding the white matter/ventricle signal regression (REG[WM, Vent] at RTP7), no connectivity showed a significant cardiac noise ratio. No connectivity-wise significant difference of the cardiac noise *R*_n_^2^ was seen in any connectivity at any sequential contrast.

For the respiration noise, the connectivity-wise analysis showed no significant *R*_n_^2^ and *R*_n_^2^differences between pipelines in any connectivity.

#### 3.3.3 RTP effects on the noise variance ratio (*R*_n_^2^) in the two-point connectivity

Figure 5C shows distributions of the mean noise variance ratio (*R*_n_^2^) explained by the motion and physiological noises for the two-point dynamic FC time-series. The plot shows the distribution of the participant mean values of all pairwise connectivity between the 264 functional ROIs. Any significant decrease or increase of the noise ratio when adding an RTP component is summarized in Table 4 (the statistical values for each comparison are shown in Supplementary Table S3). Slice-timing correction (TSHIFT) significantly reduced the mean cardiac and respiration noise ratios. Spatial smoothing (SMOOTH) increased the cardiac noise ratio. The mean cardiac noise ratio was also decreased by the regressions of motion parameters (REG[Mot]), motion derivatives (REG[dMot]), white matter/ventricle signals (REG[WM, Vent]), and RETROICOR (REG[RICOR]). The motion derivative regression also reduced the mean respiration noise ratio.

**Table 4.** Summary of the RTP noise reduction performance for the two-point connectivity time-course

| RTP | Noise |  |  |
| --- | --- | --- | --- |
|  | Motion | Cardiac | Respiration |
| VOLREG+TSHIFT | - | ○* | ○*** |
| +SMOOTH | - | Δ** | - |
| +REG[HPF] | - | - | - |
| +REG[Mot] | - | ○* | - |
| +REG[dMot] | - | ○* | ○** |
| +REG[GS] | - | - | - |
| +REG[WM, Vent] | - | ○* | - |
| +REG[RETROICOR] | - | ○*** | - |
| +REG[RVT] | - | - | - |

Connectivity-wise analysis results are shown in figures 11 and 12. For the motion noise, the connectivity-wise analysis showed no significant *R*_n_^2^ and *R*_n_^2^ differences between pipelines in any connectivity. Figure 11 shows maps of the connectivity-wise significant cardiac noise *R*_n_^2^. Without RETROICOR, a significant cardiac noise ratio was seen in the whole-brain connectivity, and the effect was large in the connectivity between the ROIs with a significant cardiac noise ratio in the voxel-wise analysis (Fig. 7). The significant cardiac noise effect disappeared after adding the RETROICOR regressor (REG[RICOR] at RTP8). Supplementary figure S4 shows connectivity-wise significant differences in the cardiac noise *R*_n_^2^ at each sequential contrast. A significant increase was seen when adding the spatial smoothing (SMOOTH) in the areas affected by the cardiac noise in the voxel-wise analysis (Figure 7). A significant decrease was seen when adding the RETROICOR (REG[RICOR] at RTP8) in the whole brain with a substantial effect in the areas affected by the cardiac noise in the voxel-wise analysis.

**Figure 11.**
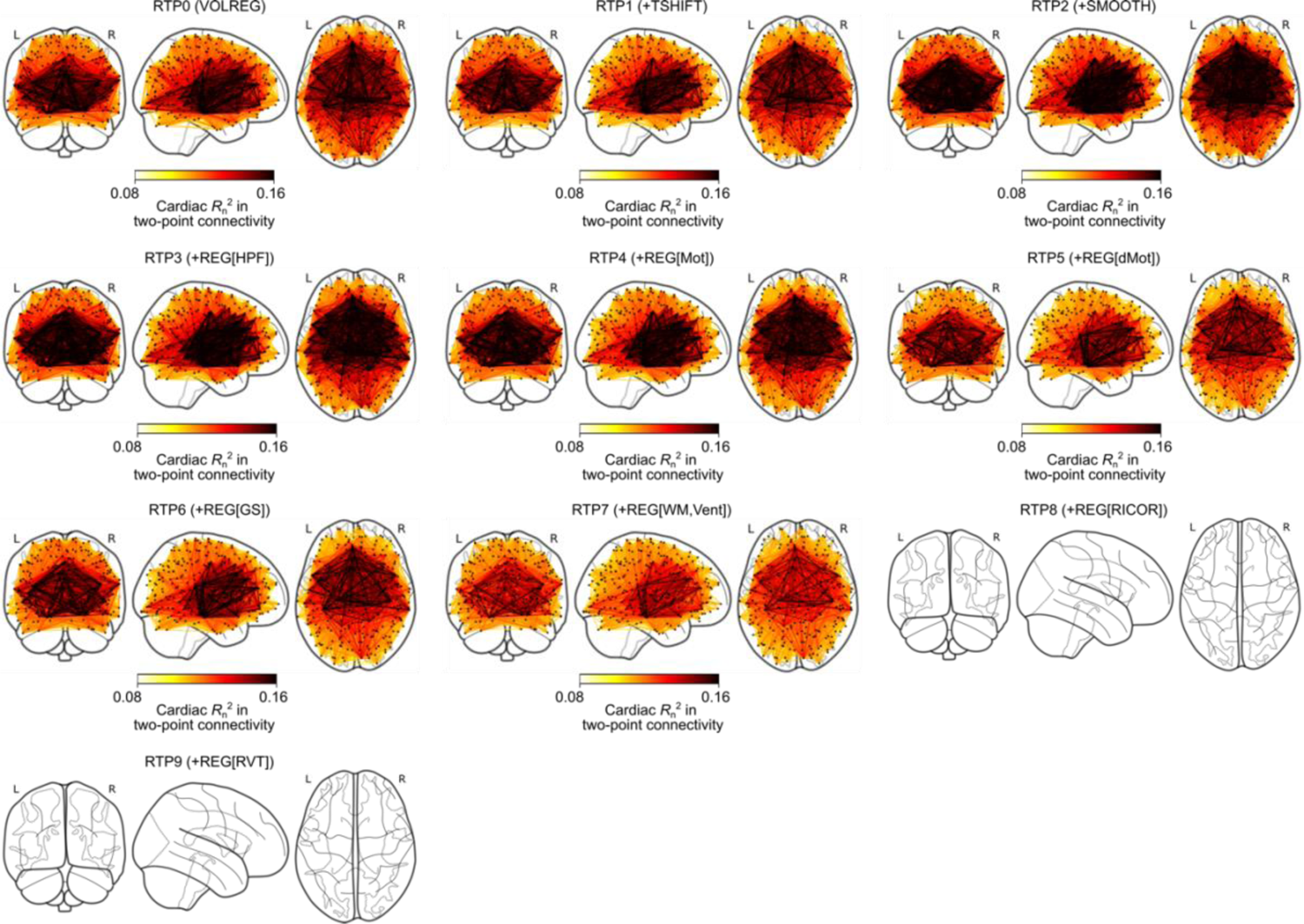
Connectivity plots of significant *R_n_*^2^ of cardiac noise in two-point connectivity time-course for each real-time processing pipeline. The plots were thresholded by FDR < 0.05 with a randomization test.

**Figure 12.**
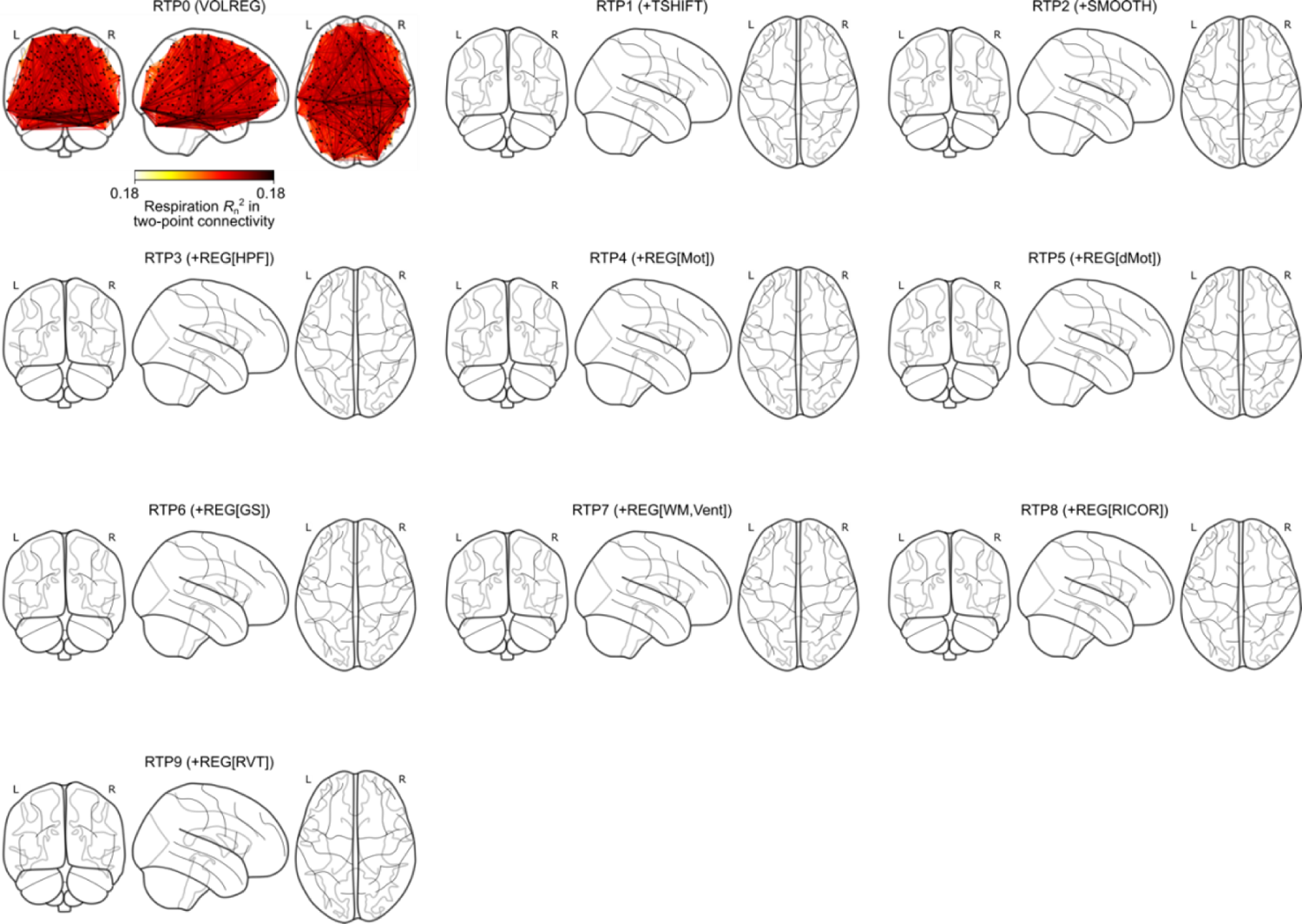
Connectivity plots of significant *R_n_* ^2^ of respiration noise in two-point connectivity time-course for each real-time processing pipeline. The plots were thresholded by FDR < 0.05 with a randomization test.

Figure 12 shows maps of the connectivity-wise significant respiration noise *R*_n_^2^. A significant respiration noise ratio was seen only for the RTP0 (only the motion correction) in the whole-brain area. No connectivity-wise significant difference of the respiration noise *R*_n_^2^ was seen in any connectivity at any sequential contrast.

#### 3.3.4 Group-level association of the mean connectivity with the motion and physiological noises

Figure 13A shows the association of the participant-wise mean sliding-window connectivity with the mean motion (frame-wise displacement), the standard deviation of heart rate, and the standard deviation of the respiration rate for RTP0 (only the motion correction) and RTP6 (added the global signal regression). Plots for all pipelines are shown in Supplementary figures S5, S6, and S7. The figures indicate that the mean connectivity was shifted to a positive value for many participants and the size of the shift was correlated with the participant’s amplitude of motion, heart rate, and respiration measures for RTP0. Adding the global signal regression (RTP6) eliminated this bias. However, the correlations (Spearman’s rank-order correlation) were still statistically significant even after the global signal regression, although the effect size (slope of the regression) became small.

**Figure 13.**
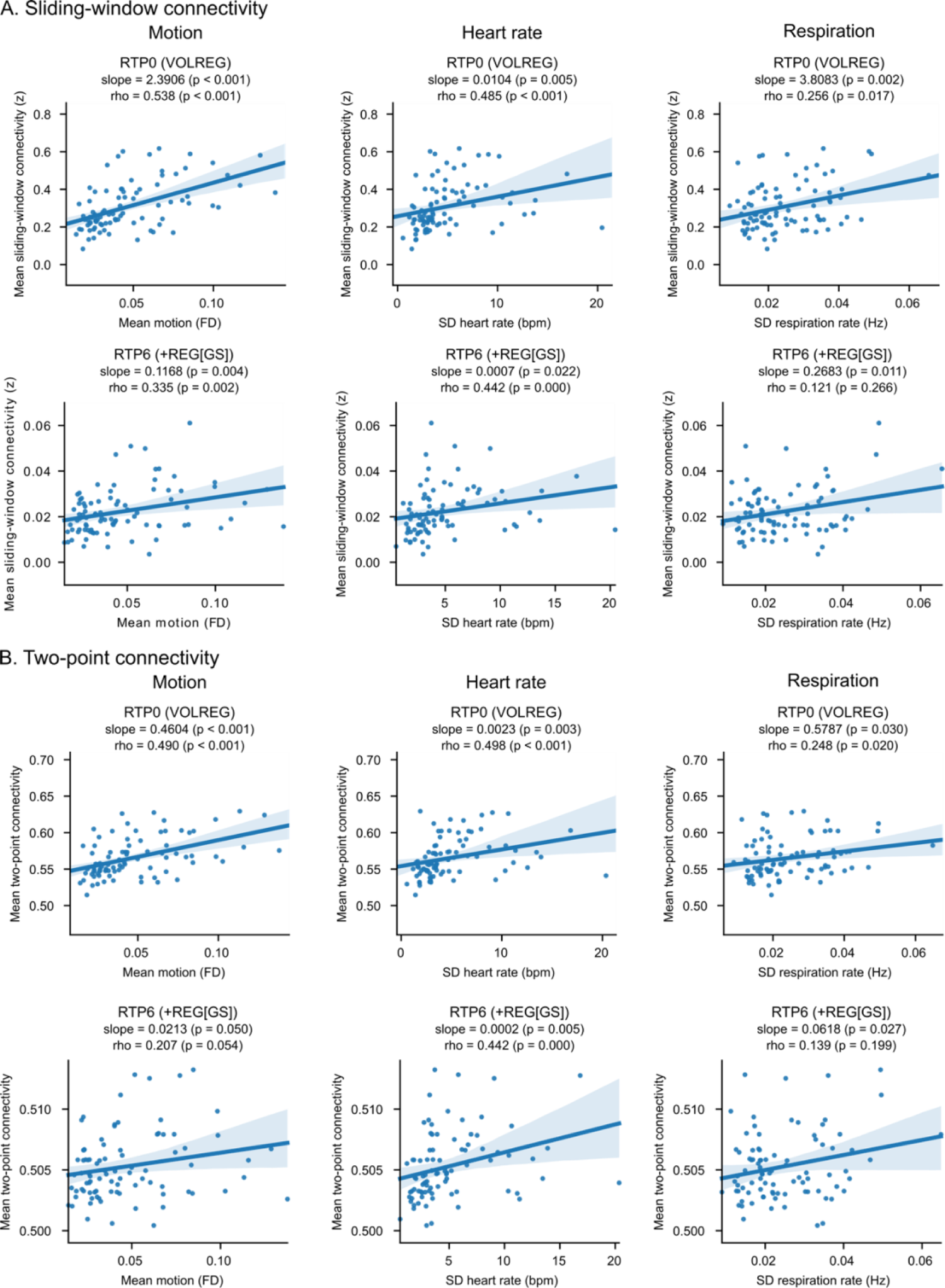
Associations between the mean connectivity (A, sliding-window [5-TR width]; B, two-point) and the mean motion (frame-wise displacement, FD), the standard deviation (SD) of heart rate and the standard deviation of respiration rate. Each point indicates a participant. The shadow around the line indicates a 95% confidence interval. The slope is a fitted coefficient of the motion in linear regression analysis, and rho is Spearman’s rank-order correlation.

Figure 13B shows the same association for the two-point connectivity. Figures for all pipelines are shown in Supplementary figures S8, S9, and S10. Like the sliding-window connectivity, the mean two-point connectivity was correlated with the participant’s amplitude of motion, heart rate, and respiration measures for RTP0. With the global signal regression (RTP6), the mean connectivity became nearly equal to 0.5 for all participants, while the associations remained statistically significant with a small effect size.

### 3.4 Noise evaluation in the real-time processed signal and signal integrity for neurofeedback task data

Figure 14A shows distributions of the noise variance ratio (*R*_n_^2^) explained by the motion and physiological noises for the left amygdala signal in the neurofeedback task run. The RTP labels with boldface indicate *R*_n_^2^ was significantly high with permutation test. The result was similar to the voxel-wise analysis of the resting-state data in the amygdala region. No significant motion *R*_n_^2^ was observed. The cardiac *R*_n_^2^ was significant in all pipelines and was significantly decreased with RETROICOR regression (RTP8). The respiration *R*_n_^2^ was significant until RTP3, and motion regression (RTP4) significantly reduced it.

**Figure 14.**
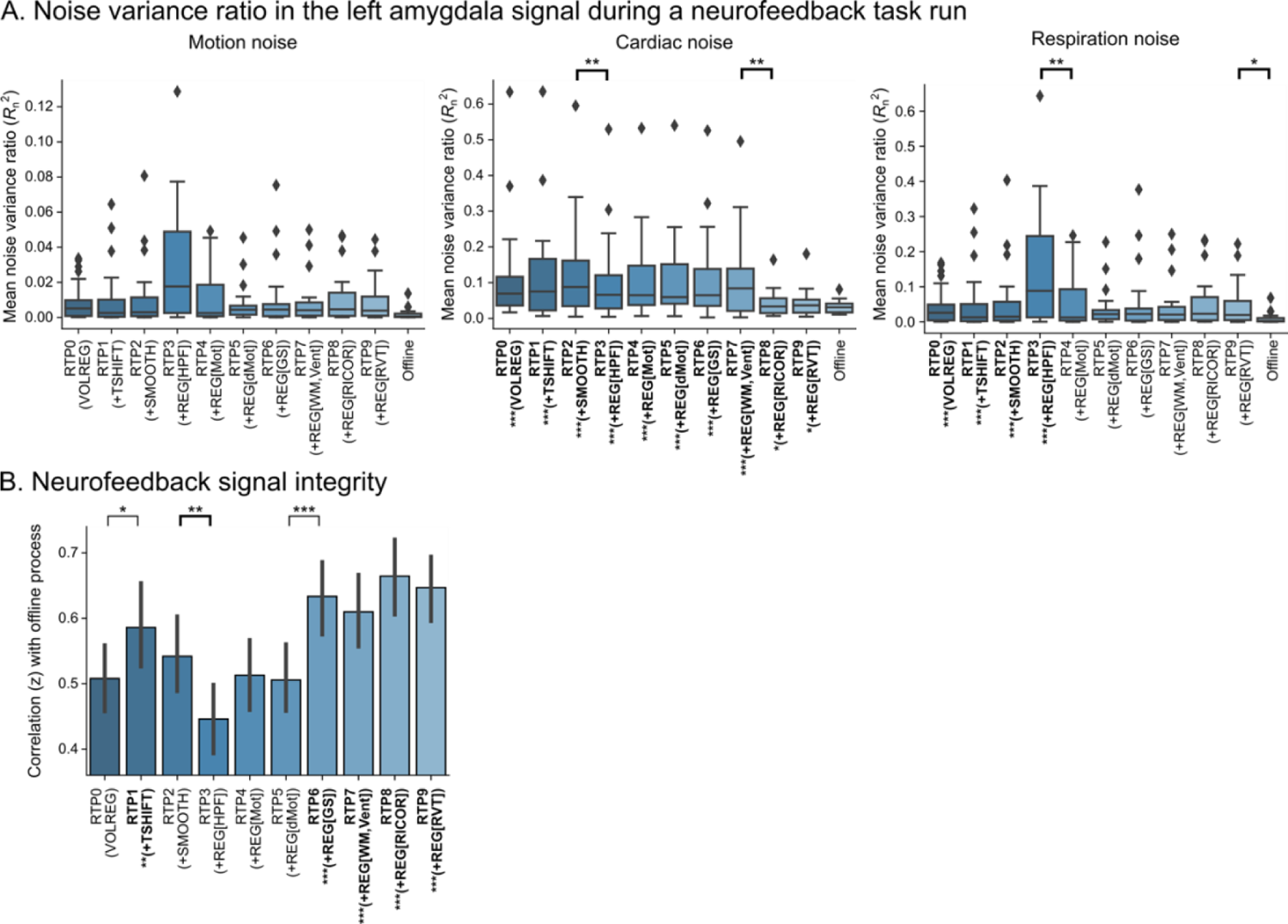
A. Box plot of the *R_n_*^2^ of noise regression on the left amygdala signal during a neurofeedback training run. The lines with starts indicate the significant difference between the pipelines; thin lines indicate increased and thick lines indicate decreased noise variance ratio. Boldface labels in the horizontal axis indicate a significant amount of *R*_n_^2^ by permutation test. B. The mean z-transformed correlation between the real-time and the offline processed left amygdala signal during a neurofeedback training run. The lines with starts indicate the significant difference between the pipelines; thin lines indicate increased and thick lines indicate decreased correlation. Boldface label in the horizontal axis indicates a significant increase in the correlation compared to RTP0. HPF, high-pass filtering regressor; Mot, motion regressor; dMot, motion derivative regressor; GS, global signal regressor; WM, Vent, white matter/ventricle mean signals regressors; RICOR, RETROIOCR regressor; RVT, respiration volume time-course regressor; *, *p* < 0.05; **, *p* < 0.01; ***, *p* < 0.001. P-values were corrected with false discovery rate.

Figure 14B shows the mean and 95% confidence interval of the signal correlation (z-transformed) between the RTP and offline-processed signals. The RTP labels with boldface indicate that the correlation was significantly higher than RTP0. For the neurofeedback task run, the left amygdala signal with RTP8 had the highest mean correlation with the offline-processed signal.

## 4 Discussion

The simulation results indicated that 1) cumulative GLM compared to incremental GLM could substantially improve the accuracy of online-formed regressors with a retrospective correction, 2) extensive RTP with the cumulative GLM could be completed in a short enough time for real-time processing, and 3) significant noise reduction was seen for many tested RTP steps, though its benefit differs among the voxel-wise and online dynamic FC signals.

### 4.1 Cumulative GLM is preferred in RTP

The high-pass filtering regression with iGLM filtered higher frequency than the designed threshold at early TRs (Fig. 2). This deficit of filtering property is because a piece of high-pass filtering regressors made for a full-length scan could fit higher frequency components than the designed threshold. cGLM avoids this problem by updating the regressors to adjust to the designed pass frequency at each length of data.

However, the difference between the GLM approaches was not large for the BOLD signal compared to the white noise signal. This is because most of the BOLD signal power is in a low-frequency range, resulting in relatively small differences in the high-frequency range. This result suggests that the iGLM drawback of high-pass-filtering may not be a substantial problem in a practical rtfMRI application. Also, the drawback of high-pass filtering regression in iGLM may be avoided by using another online frequency filtering method, such as the FIR filter [13]. However, we should remember that we must apply the same filter to the regressors if we use noise regression other than the high-pass filtering; otherwise, an artifact in the residual signal could result [41].

In an online creation of physiological noise regressors, respiration and RVT noise models were not accurate with the incremental approach (Fig. 3). To detect a low-frequency fluctuation and its phase we need a certain length of observation (at least one cycle of fluctuation). Since the cycle of respiration is usually longer than a TR, the respiration phase estimation can be made only retrospectively. Also, the RVT regressor depends on the peak detection in the respiration signal time-series, but the peak can be identified only retrospectively. Thus, the phase estimation and peak detection cannot be accurate in an online process. As the defect of an inaccurate noise regressor was accumulated with iGLM, the correlation between the online and offline regressors continued decreasing in iGLM. In contrast, cGLM could compensate for this problem by updating the regressors retrospectively at every TR.

The iGLM has been popular in rtfMRI applications thanks to its low computer memory consumption and short computation time. However, the present simulation indicates that the cumulative approach has a notable advantage when an online refinement of the regressor is required. The simulation also demonstrated that the cost of computation time for cGLM is not so high. Consequently, we should use the cumulative GLM in RTP, especially when using online-made physiological noise regressors.

### 4.2 Computation time does not limit the extensive RTP

The computation time was short enough for real-time processing in our RTP simulation system. The total processing time was less than 400 ms with GPU and less than 500 ms with CPU only (Fig. 4). The most time-consuming part of the present system was motion correction (VOLREG). We found that resampling a data volume in a registered space took long in this process. While we used a compiled C library of AFNI implementation using CPU only, a GPU implementation, like Scheinost, Hampson, Qiu, Bhawnani, Constable and Papademetris [9], might further shorten the computation time of this process.

The REGRESS took more time in later TRs than in earlier TRs since the number of samples included in the cGLM increased with time. The slope of the processing time increase was less steep with GPU than CPU only, indicating that a cumulative GLM computation with a more extended scan is possible with GPU, as far as the memory space allows. In the current simulation, we used a relatively large matrix size (128 x 128 x 34) as fMRI and the number of voxels was 149,082 (about 600 kB with single-precision float) per volume on average after applying the brain mask. This demonstrates that the data size would not limit the cumulative computation with a recent GPU’s large gigabyte memory. Even if the computational capacity is limited compared to the present simulation, we can reduce the burden by limiting the processing regions (e.g., in the gray matter or in a target region). Thus, the computation time should not limit the application of comprehensive real-time fMRI processing.

The simulation result shows that we can almost ignore the cost of computation time in RTP with current computer hardware. However, we still need to consider another cost of RTP, the number of regressors. The more regressors we use, the longer burn-in time we need before utilizing the regression in order to wait for a sufficient number of samples. Hinds, Ghosh, Thompson, Yoo, Whitfield-Gabrieli, Triantafyllou and Gabrieli [17] showed that 25 TRs were required to make a reliable regression result. Our previous investigation [10] indicated that the required TRs depended on the number of regressors and the output of real-time regression with many regressors was unreliable in the early TRs because of overfitting. The long burn-in time could cost scan time and limit experimental design in some applications of rtfMRI. Hence, we should refrain from including a regressor without a significant effect on reducing noise.

### 4.3 RTP noise reduction for the voxel-wise signal

Spatial smoothing (SMOOTH) increased the mean noise variance ratio for the cardiac and respiration noises, and high-pass filtering (REG[HPF]) increased the mean noise variance ratio for the cardiac noise. The increased noise ratio with smoothing was partly due to the spread of the noise effect in the voxels neighboring the noise-contaminated areas (e.g., Supplementary figures S2 and S3). The increased noise variance ratio is also attributable to the decreased total variance. Since the smoothing did not specifically reduce the physiological noises, the decrease of the total variance by smoothing could increase the relative ratio of the physiological noise variance. The same logic could be applied to high-pass filtering. As the BOLD signal is dominated by low-frequency components (Fig. 2B), high-pass filtering could reduce a large amount of total variance. If the cardiac noise was not included in the removed low-frequency range, the ratio of cardiac noise variance to the total variance could increase.

We consider that these increases in noise variance ratio are not problematic. These are the results of removing different kinds of noise variance, such as high-spatial-frequency noise and low-temporal-frequency signal fluctuation. Indeed, high-pass filtering regression significantly reduced the mean motion and respiration noise ratios, indicating that this regression is beneficial in removing a low-frequency noise. Also, the slice-timing correction has indicated a significant benefit in a reliable estimation of brain activity [42], and a noticeable effect of spatial smoothing on activation detection has been demonstrated [43]. Regardless of the motion and physiological noise variance ratios, we consider that TSHIFT, VOLREG, SMOOTH, and REG[HPF] should always be included in RTP because these remove known noise components other than the motion and physiological noises.

A significant reduction of the mean noise variance ratio in any of the motion and physiological noises for the voxel-wise signal was seen with regressions of high-pass filtering (REG[HPF]), motion parameters (REG[Mot]), white matter/ventricle signals (REG[WM, Vent]), and RETROICOR (REG[RICOR]). In the voxel-wise evaluation, the motion derivatives also had a significant effect on reducing the noise in many voxels (Supplementary figure S1, S2, and S3) while not significant in the whole-brain average.

The global signal regression significantly increased the cardiac noise ratio (Supplementary figure S2) and the RVT regression increased the respiration noise ratio (Supplementary figure S3) in many voxels in the voxel-wise evaluation. The increase with global signal regression might be attributable to a reduction of the total signal variance. The increase with RVT regression could be due to the inaccuracy of the online-made RVT regressor. Although the retrospective update with cGLM was better than iGLM, the online-made regressor still showed a reduced correlation with the offline one (Fig3). The error in the online-made RVT regressor might have synchronized with the respiration signal, which could recall the respiration-correlated signal fluctuation in the residual of the RTP regression. Similar recall of noise has been demonstrated by Hallquist, Hwang and Luna [41] for the high-pass filtering regression; when a high-pass filtered signal was regressed with a regressor without high-pass filtering, the residual could include the filtered low-frequency component. This harmful effect indicates that RVT regression should not be used in RTP.

Adding the white matter/ventricle signal regression (REG[WM, Vent]) after the global signal regression (REG[GS]) showed a significant reduction of the cardiac noise (RTP7-RT6 in table S1 and figure S2 in Supplementary materials), indicating that the mean white matter/ventricle signals were distinct from the global signal. This result is consistent with the evidence that gray matter had the largest effect on the global signal and the effect of the white matter and cerebrospinal fluid was small except for the partial-volume effects [44, 45]. Also, a significant cardiac noise reduction was seen when adding RETROICOR (REG[RICOR]) after adding the global signal and white matter/ventricle signal regressors (RTP8-RTP7 in table S1 and figure S2 in Supplementary materials), indicating that the global signal and the tissue-wise average signals cannot substitute for the RETROICOR regressor. Since the cardiac noise effect was concentrated on the voxels around the large blood vessels, the global and tissue-wise averages could not match such a spatially localized noise. This indicates that each of the global signal regression, white matter/ventricle signal regression, and RETROICOR regression have a distinctive contribution to noise reduction, and they cannot substitute for each other.

A potential risk of physiological noise effect on fMRI results has been indicated in the regions prone to cardiac noise [46]. The regions with significant cardiac noise (Figure 7) also overlap with the salience network area [47] and are a clinically meaningful target for intervening emotion-regulation function [48, 49]. RETORICOR regression should be vital for getting a robust neurofeedback signal from these areas. Global signal regression and mean white matter/ventricle signal regression cannot substitute for it. We also note that the cardiac noise ratio remained significant even after RETROICOR in the voxels around the large cerebral blood vessels (Figure 7, RTP8). Although the remaining noise ratio was small (< 0.1), we should be cautious of the cardiac noise when using the signal in these areas for neurofeedback.

In summary, for the online voxel-wise signal evaluation, regression with motion parameters (REG[Mot]), mean white matter/ventricle signals (REG[WM, Vent]), and RETROICOR (REG[RICOR]) significantly reduced the motion or physiological noises on average across the brain, while the effect of RTP was different across voxels. Motion derivative regressor (REG[dMot]) also reduced the noise at several voxels. While TSHIFT, VOLREG, SMOOTH, and REG[HPF] had no significant effect or increased the noise ratios due to a reduced total variance, they should be applied in RTP as they could reduce other kinds of noise. No significant contribution was found with the global signal regression (REG[GS]), and the online-made RVT regressor introduced a respiration noise artifact; thus, these regressors should not be used in RTP for the voxel-wise signals.

### 4.4 RTP noise reduction for the sliding-window connectivity

A significant reduction of the mean noise variance ratio in any of the motion and physiological noises was seen with regressions of motion derivatives (REG[dMot]), global signal (REG[GS]), and RETROICOR (REG[RICOR]) for the sliding-window dynamic FC. Global signal regression had a significant effect on reducing the motion noise variance ratio. This is consistent with the report by Parkes, Fulcher, Yucel and Fornito [50], showing the benefit of global signal regression in reducing motion noise for an offline resting-state fMRI analysis. The significant reduction of the cardiac noise with RETROICOR indicates that the global signal and the tissue-wise average signals cannot substitute for the RETROICOR regressor, which was the same as in the voxel-wise signal.

With the connectivity-wise evaluation, significant motion noise was seen in the whole-brain area, that was disappeared after adding the motion derivative regression (RTP5 in Fig. 9). A significant cardiac noise ratio was localized in the deep brain regions, including the inferior occipital, medial to anterior temporal, and the anterior cingulate areas (Fig. 10), which overlap with large cerebral blood vessels. The connectivity in these areas was often implicated in emotion-regulation functions [48, 49]. The present result demonstrates the importance of extensive real-time denoising when we target these connectivities in a neurofeedback application.

Regarding the group-level association (Fig. 13A and Supplementary figures S5, S6, and S7), only the global signal regression could eliminate the noise-associated bias of the mean connectivity, consistent with the observation by Weiss, Zamoscik, Schmidt, Halli, Kirsch and Gerchen [18]. We will discuss the effect of global signal regression in RTP in a later independent section.

In summary, for the online sliding-window dynamic FC evaluation, regression with motion derivative (REG[dMot]), global signal (REG[GS]), and RETROICOR (REG[RICOR]) significantly reduced the motion or physiological noises on average across the brain, while the effect was different across regions. REG[GS] also removed the bias of average connectivity across time and brain regions. While TSHIFT, VOLREG, SMOOTH, and REG[HPF] had no significant effect, we think that they should be applied in RTP as they could reduce other kinds of noise, as we discussed above. Although no significant contribution was found with the motion parameter regression (REG[Mot]), when we removed this regressor from the pipeline, we observed a significant increase in the mean motion noise variance ratio compared to the best-performed pipeline (RTP8; *t* = 3.114, *p* = 0.002). This suggests that global signal regression cannot substitute for the motion parameter regression and that these regressors had distinctive contributions to motion noise reduction. Mean white matter/ventricle signals (REG[WM, Vent]) and RVT regressions had no significant contribution to the noise reduction.

### 4.5 RTP noise reduction for the two-point connectivity

Overall, the mean noise variance ratio for the two-point connectivity was less than that for the 5-TR sliding-window connectivity (the vertical axis scale in Fig. 5C is smaller than that in Figs. 5B), which could be due to the narrow window width preventing from spreading the noise effect. A significant reduction of the mean noise variance ratio in any of the motion and physiological noises was seen with TSHIFT and regressions of motion parameters (REG[Mot]), motion derivatives (REG[dMot]), white matter/ventricle signal (REG[WM, Vent]), and RETROICOR (REG[RICOR]). With the connectivity-wise evaluation, no significant effect of motion was seen in any connectivity. This is consistent with our previous investigation, demonstrating that the two-point method was less prone to motion than the sliding-window methods [32].

The two-point connectivity showed more statistically significant cardiac noise than in the sliding-window connectivity. The connectivity-wise evaluation showed a significant cardiac noise variance ratio across the whole brain (Fig. 11). Because the two-point connectivity evaluates the change direction between the two consecutive time points, it could be more sensitive to a high-temporal-frequency noise than the sliding-window method. The cardiac noise effect was still significant after REG[GS] and REG[WM, Vent], but disappeared after adding the RETROICOR regression (Fig. 11, RTP8). This demonstrates that the RETROICOR benefit cannot be replaced by global signal and tissue-wise average signals. Significant respiration noise was removed by adding slice-timing correction (TSHIFT, Fig. 12), suggesting that the two-point connectivity is sensitive to a small temporal shift by the slice-timing difference. The slice-timing difference has been known to make a deviation from true connectivity [51].

Regarding the group-level association (Fig. 13B, Supplementary figures S8, S9, S10), only the global signal regression could eliminate the noise-associated bias of the mean connectivity. This was the same as for the sliding-window connectivity and consistent with the observation by Weiss, Zamoscik, Schmidt, Halli, Kirsch and Gerchen [18].

In summary, for the online two-point dynamic FC evaluation, slice-timing correction (TSHIFT) and regression with motion parameters (REG[Mot]), motion derivatives (REG[dMot]), white matter/ventricle signal (REG[WM, Vent]), and RETROICOR (REG[RICOR]) significantly reduced the physiological noises on average across the brain, while the effect was different across regions. Global signal regression (REG[GS]) removed the bias of average connectivity across time and brain regions. While VOLREG, SMOOTH, and REG[HPF] had no significant effect or increased the noise ratios due to a reduced total variance, we think that they should be applied in RTP as they could reduce other kinds of noise. Regression with mean white matter/ventricle signals (REG[WM, Vent]) and RVT had no significant contribution to the noise reduction.

### 4.6 Global signal regression in RTP

The global signal regression had a significant effect on reducing the motion noise in the 5-TR sliding-window dynamic FC (Table 3). It also had a notable effect on reducing the average connectivity bias across time and brain regions in the group-level analysis for both the sliding-window and the two-point connectivity (Fig. 13 and Supplementary figures S5 to S10). This result is consistent with the reports for offline resting-state functional connectivity analysis [22] and the neurofeedback study with dynamic FC [18]. The neurofeedback study showed that global signal regression could remove the correlation between the participant-wise average dynamic FC and respiration measures. They also indicated that regression with the physiological noise models, including RETROICOR, could not remove this bias.

However, the present results showed that the RETROICOR regression had an additional benefit in reducing the cardiac noise after the global signal regression in the individual participant’s signal time-course. Also, even with the global signal regression, excluding the motion parameters significantly increased the motion noise variance ratio for the sliding-window connectivity (Section 4.4), indicating that the distinctive effect of motion parameter regression from the global signal regression. Thus, the global signal regression is not the all-around solution for noise reduction in functional connectivity, and the motion parameter and RETROICOR regressors significantly contributed to noise reduction even when accompanied by the global signal regression.

The discrepancy between the group-level association and the individual noise regression analysis suggests that the global signal regression removed a signal component distinct from the motion and RETROICOR regressors. The noise component inducing the average connectivity bias and being removed by the global signal regression might be a BOLD signal fluctuation having a loose temporal association with the noise time-course. Indeed, Power, Schlaggar and Petersen [22] reported that the increase of the functional connectivity by motion could be seen even in 8 to 10 s delays, which might be due to a spin-history noise [21].

The global signal regression is the most controversial processing used in resting-state fMRI. Many studies have investigated the source of the global signal and its association with noise and functional neural activations [45, 52–56]. The side effects of the global signal regression, such as an artifactual negative correlation, have also been demonstrated [57–59]. Although we refrain from further considering the source of the global signal, which is out of the scope of the present study, we consider a practical suggestion in using the global signal regression for a rtfMRI application.

In a rtfMRI application, a signal fluctuation with a tight temporal association with a motion or physiological noise should be more apparent for a participant, and such a noise could increase the risk of inducing spurious training with noise regulation. Thus, regardless of using global signal regression, we should use the motion and RETROICOR regressions in RTP. When we use a voxel-wise or an ROI-based neurofeedback signal, the present result indicates that the global signal regression is not needed in RTP. The global average signal was not useful in removing a voxel-wise noise.

When we use dynamic FC as a neurofeedback signal, we should include the global signal regression as it has a notable benefit in reducing the bias in the average connectivity across time. However, we should also be careful about a possible artifact due to the global signal regression. Once we determine the target connectivity, we should perform an RTP simulation analysis, like Ramot and Gonzalez-Castillo [60] and Misaki, Tsuchiyagaito, Al Zoubi, Paulus, Bodurka and Tulsa [32], to confirm that the global signal regression does not make a spurious shift of the connectivity neurofeedback signal. If a notable artifact is seen, we might have to refrain from using the global signal regression in RTP and admit the reduced signal-to-noise ratio in the neurofeedback signal or consider another target.

The group-level association for the mean connectivity with the motion and physiological noises remained statistically significant even after the global signal regression (Figs. 8 and 9). However, we consider that this statistical significance would not matter in the practice of neurofeedback training because the effect size of the noise (slope of the regression line) was minimal after the global signal regression. In the neurofeedback context, we can ignore the noise effect if its effect size was much smaller than that by self-regulation.

### 4.7 RTP noise reduction for the neurofeedback target signal

To confirm that the simulation results for the resting-state data could be applied to a neurofeedback task run, we performed the same simulation analysis for the neurofeedback task data [26]. The result was parallel to the resting-state data’s voxel-wise analysis. In this neurofeedback experiment, participants were trained to increase the left amygdala (LA) activation by recalling a positive autobiographical memory. Hence, the LA signal should include a self-regulated change so that the noise variance ratios could be less dominant than in a resting state. However, we still observed significant noise variance ratios. While the motion noise was not significant in any pipelines, the respiration noise was significant before adding the motion parameter regression, and the cardiac noise was significant in all RTPs with a significant reduction by RETROICOR regression (Fig. 14A). This was consistent with the resting-state data analysis result in Fig. 7, showing that the cardiac noise ratio was significantly reduced with RETROICOR, while it was still significant in the amygdala region. The signal integrity analysis (Fig. 14B) also demonstrated that the signal correlation between RTP and the offline process improved with the inclusion of additional noise regressors.

We should note that the noise amount and the RTP effect in a neurofeedback signal could vary depending on the target region, the applied regulation strategy, and the individual’s self-regulation performance. Thus, we used resting-state data to evaluate the general effect of noise and RTPs. Nevertheless, the results for the neurofeedback task data demonstrated that the noise effect could be significant even with a task-related signal change and RTP noise reduction is beneficial in the neurofeedback application.

### 4.8 Limitations

Several limitations of the current study should be acknowledged. Since we used similar regressors in RTP and the noise regression analysis, it may look trivial that including these regressors in RTP could reduce the noises. However, we should point out that the RTP regressors and offline-made noise regressors used in the noise regression analysis cannot be identical. Online real-time processing can use limited samples of the current and past time points. Indeed, we found the deficit of the RVT regression in RTP. Furthermore, the physiological noise variance extracted in the noise regression analysis was not specific to the RETROICOR model. The RETROICOR regressor has a general form of the Fourier basis set that can fit any temporal fluctuation synchronized with the cardiac and respiration fluctuations. We also included the heart rate and respiration variance convolved with their hemodynamic response functions in the noise regression analysis. Thus, the benefit of RETROICOR regressors in RTP is never a trivial result.

The present result is limited to the noise measures used for the evaluation. We used the motion (frame-wise displacement), heart rate, respiration variance, RETORICOR, and RVT in the noise evaluation. Although these are major noise sources for the BOLD signal, there are yet other sources that could have a significant effect on the signal change. For example, Power, Lynch, Dubin, Silver, Martin and Jones [34] indicated that deep breath had a large effect on the fMRI signal changes and that that cannot be captured by the respiration envelope, respiration variance, and RVT measures. In a rtfMRI application, complete noise-cleaning may be too costly since using more RTP regressors increases the initial burn-in time further. However, at the offline analysis in evaluating the neurofeedback effect, we should examine the noise effect that has not been removed at RTP. The apparent training effect could result from a change in motion or physiological noise pattern [18].

Using resting-state data in the simulation might also limit the implication of the present results. When there was a task-induced signal change, the noise variance ratio may be less significant than in the present results. However, a substantial signal change is not always guaranteed, especially in the early phase of the training when self-regulation ability is not high. Even if a participant could regulate the brain activation, a task-induced fMRI signal change is often smaller than the noise variance; temporal contrast to noise ratio is usually lower than 1.0 [61, 62]. For detecting such a low contrast signal change, noise reduction should be applied as much as possible. Indeed, the simulation for the neurofeedback task data demonstrated that the noise variance ratio was still significant with a self-regulation task. Furthermore, if a noise-contaminated neurofeedback signal was used, the training could depend on the regulation of noise-associated motion or physiological states. Thus, the caveat of the present results could be applied to general neurofeedback training.

While we presented beneficial RTP components for the voxel-wise and dynamic FC signals on average across the brain regions, the optimal combination with minimum regressors will depend on the target regions. For example, the cardiac noise effect concentrated on the areas overlapping with large cerebral blood vessels (Fig. 7), which was removed by RETROICOR regression. If the target is out of these regions, RETORICOR might not be needed. Although the present results help consider which kinds of noise could affect which brain areas and what RTP helps reduce it, the optimal RTP pipeline needs to be designed for each RTP application with a simulation analysis [60, 63].

Regarding the sliding-window dynamic FC, the RTP effect could be different depending on window width. The previous study [32] indicated that the correlation between motion and dynamic FC increased with the window width so that a narrow window was more robust to noise. Also, a narrow window could be preferred in a neurofeedback application to provide a timely feedback signal; a wide-window dynamic FC reflects less current brain activation and depends more on the long history of past brain states. However, several studies with an offline analysis have shown that a wide window is required to evaluate dynamic FC robustly [64], and that even with a wide window (e.g., 2 min), dynamic FC is challenging to detect with a single session measurement [65]. Thus, in choosing the window width for a dynamic FC neurofeedback, we need to consider the trade-off between the robust evaluation of brain state and timely and less noisy feedback. While the optimal choice could be different between individual applications, we consider that a narrow window can be approved in a neurofeedback application. We should also note that a TR-wise estimate of regional brain activation is not so robust compared to an offline evaluation with many time points, and many studies have proven that even such a fragile estimate helped participants learn brain self-regulation. Thus, even a less robust estimate of brain activation could be useful in training participants to self-regulate their brain state. Indeed, studies with the two-point dynamic FC neurofeedback, which is one extreme window width focusing on feedback timeliness, demonstrated successful training in self-regulating functional connectivity [30, 66].

While using a control ROI or control connectivity was not considered in the present study, it could be an economical noise reduction regressor. However, we should remember that the control selection should be specific to the target region and is not a trivial task. The present simulation demonstrated that the global signal and the white matter/ventricle average signals could not be a substitute for the RETROICOR, suggesting that just taking the average of the regions apart from the target would not be enough to suppress the noise. Also, if the selected control area had a unique functional response, subtracting the control signal from the target could induce an artifactual signal fluctuation. The self-regulation task usually invites brain activations in many regions, not limited to the target [67–71]. Thus, we must be very cautious in choosing an appropriate control signal, which needs an extensive independent evaluation to optimize noise reduction performance.

## 5 Conclusions

The present study demonstrated that extensive real-time noise reduction had a minimal cost of computation time with significant benefit in reducing motion and physiological noises for rtfMRI. We believe our comprehensive evaluation will promote a widespread utilization of extensive real-time noise reduction for neurofeedback applications. The high-quality real-time estimate of brain activity would pave the way for robust and replicable neurofeedback training, as well as the urgently needed development of novel and robust interventions for several mental and neurological disorders, as well as a potential application in presurgical mapping.

## Acknowledgement

This research was supported by the Laureate Institute for Brain Research (LIBR) and the William K. Warren Foundation, and in part by the National Institute of General Medical Sciences, National Institutes of Health 1P20GM121312 award. The simulation samples were provided by NIMH/NIH grant R01 MH098099. The funding agencies were not involved in the design of experiment, data collection and analysis, interpretation of results and preparation and submission of the manuscript.

## Notes

### Competing Interest Statement

The authors have declared no competing interest.

## References

1. Cox R W, Jesmanowicz A and Hyde J S 1995 Real-time functional magnetic resonance imaging Magnetic Resonance in Medicine 33 230–6

2. Weiskopf N, Sitaram R, Josephs O, Veit R, Scharnowski F, Goebel R, Birbaumer N, Deichmann R and Mathiak K 2007 Real-time functional magnetic resonance imaging: methods and applications Magnetic resonance imaging 25 989-1003

3. Goebel R, Zilverstand A and Sorger B 2010 Real-time fMRI-based brain-computer interfacing for neurofeedback therapy and compensation of lost motor functions Imaging in Medicine 2 407–15

4. Sulzer J, Haller S, Scharnowski F, Weiskopf N, Birbaumer N, Blefari M L, Bruehl A B, Cohen L G, DeCharms R C, Gassert R, Goebel R, Herwig U, LaConte S, Linden D, Luft A, Seifritz E and Sitaram R 2013 Real-time fMRI neurofeedback: progress and challenges Neuroimage 76 386-99

5. Weiskopf N, Scharnowski F, Veit R, Goebel R, Birbaumer N and Mathiak K 2004 Self-regulation of local brain activity using real-time functional magnetic resonance imaging (fMRI) Journal of Physiology Paris 98 357-73

6. Heunis S, Lamerichs R, Zinger S, Caballero-Gaudes C, Jansen J F A, Aldenkamp B and Breeuwer M 2020 Quality and denoising in real-time functional magnetic resonance imaging neurofeedback: A methods review Hum Brain Mapp 41 3439-67

7. Thibault R T, MacPherson A, Lifshitz M, Roth R R and Raz A 2018 Neurofeedback with fMRI: A critical systematic review Neuroimage 172 786-807

8. Bagarinao E, Matsuo K, Nakai T and Sato S 2003 Estimation of general linear model coefficients for real-time application Neuroimage 19 422-9

9. Scheinost D, Hampson M, Qiu M, Bhawnani J, Constable R T and Papademetris X 2013 A graphics processing unit accelerated motion correction algorithm and modular system for real-time fMRI Neuroinformatics 11 291-300

10. Misaki M, Barzigar N, Zotev V, Phillips R, Cheng S and Bodurka J 2015 Real-time fMRI processing with physiological noise correction – Comparison with off-line analysis Journal of Neuroscience Methods 256 117-21

11. Goebel R 2012 BrainVoyager--past, present, future Neuroimage 62 748–56

12. Heunis S, Besseling R, Lamerichs R, de Louw A, Breeuwer M, Aldenkamp B and Bergmans J 2018 Neu3CA-RT: A framework for real-time fMRI analysis Psychiatry Research: Neuroimaging 282 90-102

13. Koush Y, Ashburner J, Prilepin E, Sladky R, Zeidman P, Bibikov S, Scharnowski F, Nikonorov A and De Ville D V 2017 OpenNFT: An open-source Python/Matlab framework for real-time fMRI neurofeedback training based on activity, connectivity and multivariate pattern analysis Neuroimage 156 489-503

14. Stoeckel L E, Garrison K A, Ghosh S, Wighton P, Hanlon C A, Gilman J M, Greer S, Turk-Browne N B, deBettencourt M T, Scheinost D, Craddock C, Thompson T, Calderon V, Bauer C C, George M, Breiter H C, Whitfield-Gabrieli S, Gabrieli J D, LaConte S M, Hirshberg L, Brewer J A, Hampson M, Van Der Kouwe A, Mackey S and Evins A E 2014 Optimizing real time fMRI neurofeedback for therapeutic discovery and development Neuroimage Clin 5 245-55

15. Ros T, Enriquez-Geppert S, Zotev V, Young K D, Wood G, Whitfield-Gabrieli S, Wan F, Vuilleumier P, Vialatte F, Van De Ville D, Todder D, Surmeli T, Sulzer J S, Strehl U, Sterman M B, Steiner N J, Sorger B, Soekadar S R, Sitaram R, Sherlin L H, Schonenberg M, Scharnowski F, Schabus M, Rubia K, Rosa A, Reiner M, Pineda J A, Paret C, Ossadtchi A, Nicholson A A, Nan W, Minguez J, Micoulaud-Franchi J A, Mehler D M A, Luhrs M, Lubar J, Lotte F, Linden D E J, Lewis-Peacock J A, Lebedev M A, Lanius R A, Kubler A, Kranczioch C, Koush Y, Konicar L, Kohl S H, Kober S E, Klados M A, Jeunet C, Janssen T W P, Huster R J, Hoedlmoser K, Hirshberg L M, Heunis S, Hendler T, Hampson M, Guggisberg A G, Guggenberger R, Gruzelier J H, Gobel R W, Gninenko N, Gharabaghi A, Frewen P, Fovet T, Fernandez T, Escolano C, Ehlis A C, Drechsler R, Christopher deCharms R, Debener S, De Ridder D, Davelaar E J, Congedo M, Cavazza M, Breteler M H M, Brandeis D, Bodurka J, Birbaumer N, Bazanova O M, Barth B, Bamidis P D, Auer T, Arns M and Thibault R T 2020 Consensus on the reporting and experimental design of clinical and cognitive-behavioural neurofeedback studies (CRED-nf checklist) Brain 143 1674-85

16. Kopel R, Sladky R, Laub P, Koush Y, Robineau F, Hutton C, Weiskopf N, Vuilleumier P, Van De Ville D and Scharnowski F 2019 No time for drifting: Comparing performance and applicability of signal detrending algorithms for real-time fMRI Neuroimage 191 421-9

17. Hinds O, Ghosh S, Thompson T W, Yoo J J, Whitfield-Gabrieli S, Triantafyllou C and Gabrieli J D E 2011 Computing moment-to-moment BOLD activation for real-time neurofeedback NeuroImage 54 361-8

18. Weiss F, Zamoscik V, Schmidt S N L, Halli P, Kirsch P and Gerchen M F 2020 Just a very expensive breathing training? Risk of respiratory artefacts in functional connectivity-based real-time fMRI neurofeedback Neuroimage 210 116580

19. Glover G H, Li T Q and Ress D 2000 Image-based method for retrospective correction of physiological motion effects in fMRI: RETROICOR Magn Reson Med 44 162–7

20. Birn R M, Smith M A, Jones T B and Bandettini P A 2008 The respiration response function: the temporal dynamics of fMRI signal fluctuations related to changes in respiration NeuroImage 40 644-54

21. Friston K J, Williams S, Howard R, Frackowiak R S and Turner R 1996 Movement-related effects in fMRI time-series Magn Reson Med 35 346-55

22. Power J D, Schlaggar B L and Petersen S E 2015 Recent progress and outstanding issues in motion correction in resting state fMRI Neuroimage 105 536–51

23. Nikolaou F, Orphanidou C, Papakyriakou P, Murphy K, Wise R G and Mitsis G D 2016 Spontaneous physiological variability modulates dynamic functional connectivity in resting-state functional magnetic resonance imaging Philos Trans A Math Phys Eng Sci 374 20150183

24. Kruger G and Glover G H 2001 Physiological noise in oxygenation-sensitive magnetic resonance imaging Magn Reson Med 46 631–7

25. Birn R M, Diamond J B, Smith M A and Bandettini P A 2006 Separating respiratory-variation-related fluctuations from neuronal-activity-related fluctuations in fMRI NeuroImage 31 1536-48

26. Young K D, Zotev V, Phillips R, Misaki M, Yuan H, Drevets W C and Bodurka J 2014 Real-time FMRI neurofeedback training of amygdala activity in patients with major depressive disorder PLoS One 9 e88785

27. Misaki M, Suzuki H, Savitz J, Drevets W C and Bodurka J 2016 Individual Variations in Nucleus Accumbens Responses Associated with Major Depressive Disorder Symptoms Scientific reports 6 21227

28. Power J D, Cohen A L, Nelson S M, Wig G S, Barnes K A, Church J A, Vogel A C, Laumann T O, Miezin F M, Schlaggar B L and Petersen S E 2011 Functional network organization of the human brain Neuron 72 665-78

29. Gembris D, Taylor J G, Schor S, Frings W, Suter D and Posse S 2000 Functional magnetic resonance imaging in real time (FIRE): Sliding-window correlation analysis and reference-vector optimization Magnetic Resonance in Medicine 43 259-68

30. Ramot M, Kimmich S, Gonzalez-Castillo J, Roopchansingh V, Popal H, White E, Gotts S J and Martin A 2017 Direct modulation of aberrant brain network connectivity through real-time NeuroFeedback Elife 6 e28974

31. Mokhtari F, Akhlaghi M I, Simpson S L, Wu G and Laurienti P J 2019 Sliding window correlation analysis: Modulating window shape for dynamic brain connectivity in resting state Neuroimage 189 655-66

32. Misaki M, Tsuchiyagaito A, Al Zoubi O, Paulus M, Bodurka J and Tulsa I 2020 Connectome-wide search for functional connectivity locus associated with pathological rumination as a target for real-time fMRI neurofeedback intervention Neuroimage Clin 26 102244

33. Chang C, Cunningham J P and Glover G H 2009 Influence of heart rate on the BOLD signal: the cardiac response function NeuroImage 44 857–69

34. Power J D, Lynch C J, Dubin M J, Silver B M, Martin A and Jones R M 2020 Characteristics of respiratory measures in young adults scanned at rest, including systematic changes and “missed” deep breaths Neuroimage 204 116234

35. Bates D, Maechler M, Bolker B and Walker S 2015 Fitting Linear Mixed-Effects Models Using lme4 Journal of Statistical Software 67 1–48

36. Kuznetsova A, Brockhoff P B, B. R H and Christensen G E 2017 lmerTest Package: Tests in Linear Mixed Effects Models Journal of Statistical Software 82 1–26

37. R Core Team 2018 R: A Language and Environment for Statistical Computing. R Foundation for Statistical Computing, Vienna, Austria. URL https://www.R-project.org/)

38. Lenth R V 2016 Least-squares means: the R package lsmeans J Stat Softw 69 1-33

39. Liegeois R, Laumann T O, Snyder A Z, Zhou J and Yeo B T T 2017 Interpreting temporal fluctuations in resting-state functional connectivity MRI Neuroimage 163 437-55

40. Prichard D and Theiler J 1994 Generating surrogate data for time series with several simultaneously measured variables Physical review letters 73 951–4

41. Hallquist M N, Hwang K and Luna B 2013 The nuisance of nuisance regression: spectral misspecification in a common approach to resting-state fMRI preprocessing reintroduces noise and obscures functional connectivity Neuroimage 82 208–25

42. Sladky R, Friston K J, Trostl J, Cunnington R, Moser E and Windischberger C 2011 Slice-timing effects and their correction in functional MRI Neuroimage 58 588-94

43. Mikl M, Marecek R, Hlustik P, Pavlicova M, Drastich A, Chlebus P, Brazdil M and Krupa P 2008 Effects of spatial smoothing on fMRI group inferences Magn Reson Imaging 26 490-503

44. Power J D, Mitra A, Laumann T O, Snyder A Z, Schlaggar B L and Petersen S E 2014 Methods to detect, characterize, and remove motion artifact in resting state fMRI Neuroimage 84 320-41

45. Power J D, Plitt M, Laumann T O and Martin A 2017 Sources and implications of whole-brain fMRI signals in humans Neuroimage 146 609-25

46. Boubela R N, Kalcher K, Huf W, Seidel E M, Derntl B, Pezawas L, Nasel C and Moser E 2015 fMRI measurements of amygdala activation are confounded by stimulus correlated signal fluctuation in nearby veins draining distant brain regions Scientific reports 5 10499

47. Seeley W W, Menon V, Schatzberg A F, Keller J, Glover G H, Kenna H, Reiss A L and Greicius M D 2007 Dissociable intrinsic connectivity networks for salience processing and executive control J Neurosci 27 2349-56

48. Kral T R A, Schuyler B S, Mumford J A, Rosenkranz M A, Lutz A and Davidson R J 2018 Impact of short-and long-term mindfulness meditation training on amygdala reactivity to emotional stimuli NeuroImage 181 301-13

49. Silvers J A, Insel C, Powers A, Franz P, Helion C, Martin R E, Weber J, Mischel W, Casey B J and Ochsner K N 2017 vlPFC-vmPFC-Amygdala Interactions Underlie Age-Related Differences in Cognitive Regulation of Emotion Cereb Cortex 27 3502-14

50. Parkes L, Fulcher B, Yucel M and Fornito A 2018 An evaluation of the efficacy, reliability, and sensitivity of motion correction strategies for resting-state functional MRI Neuroimage 171 415-36

51. Kiebel S J, Kloppel S, Weiskopf N and Friston K J 2007 Dynamic causal modeling: a generative model of slice timing in fMRI Neuroimage 34 1487-96

52. Aquino K M, Fulcher B D, Parkes L, Sabaroedin K and Fornito A 2020 Identifying and removing widespread signal deflections from fMRI data: Rethinking the global signal regression problem Neuroimage 212 116614

53. Li J, Bolt T, Bzdok D, Nomi J S, Yeo B T T, Spreng R N and Uddin L Q 2019 Topography and behavioral relevance of the global signal in the human brain Scientific reports 9 14286

54. Li J, Kong R, Liegeois R, Orban C, Tan Y, Sun N, Holmes A J, Sabuncu M R, Ge T and Yeo B T T 2019 Global signal regression strengthens association between resting-state functional connectivity and behavior Neuroimage 196 126-41

55. Liu T T, Nalci A and Falahpour M 2017 The global signal in fMRI: Nuisance or Information? Neuroimage 150 213–29

56. Murphy K and Fox M D 2017 Towards a consensus regarding global signal regression for resting state functional connectivity MRI Neuroimage 154 169–73

57. Murphy K, Birn R M, Handwerker D A, Jones T B and Bandettini P A 2009 The impact of global signal regression on resting state correlations: are anti-correlated networks introduced? Neuroimage 44 893–905

58. Saad Z S, Gotts S J, Murphy K, Chen G, Jo H J, Martin A and Cox R W 2012 Trouble at rest: how correlation patterns and group differences become distorted after global signal regression Brain connectivity 2 25-32

59. Fox M D, Zhang D, Snyder A Z and Raichle M E 2009 The global signal and observed anticorrelated resting state brain networks *Department of Psychology and Center for Perceptual Systems*, The University of Texas at Austin, 108 E. Dean Keeton, 1 University Station A8000, Austin, TX 78712-0187, USA. 101 3270-83

60. Ramot M and Gonzalez-Castillo J 2018 A framework for offline evaluation and optimization of real-time algorithms for use in neurofeedback, demonstrated on an instantaneous proxy for correlations Neuroimage 188 322–34

61. Koush Y, Zvyagintsev M, Dyck M, Mathiak K A and Mathiak K 2012 Signal quality and Bayesian signal processing in neurofeedback based on real-time fMRI Neuroimage 59 478-89

62. Geissler A, Gartus A, Foki T, Tahamtan A R, Beisteiner R and Barth M 2007 Contrast-to-noise ratio (CNR) as a quality parameter in fMRI J Magn Reson Imaging 25 1263-70

63. Misaki M, Tsuchiyagaito A, Al Zoubi O, Paulus M and Bodurka J 2020 Connectome-wide search for functional connectivity locus associated with pathological rumination as a target for real-time fMRI neurofeedback intervention NeuroImage: Clinical 26 102244

64. Wilson R S, Mayhew S D, Rollings D T, Goldstone A, Przezdzik I, Arvanitis T N and Bagshaw A P 2015 Influence of epoch length on measurement of dynamic functional connectivity in wakefulness and behavioural validation in sleep Neuroimage 112 169-79

65. Hindriks R, Adhikari M H, Murayama Y, Ganzetti M, Mantini D, Logothetis N K and Deco G 2016 Can sliding-window correlations reveal dynamic functional connectivity in resting-state fMRI? Neuroimage 127 242–56

66. Tsuchiyagaito A, Misaki M, Zoubi O A, Tulsa I, Paulus M and Bodurka J 2020 Prevent breaking bad: A proof of concept study of rebalancing the brain’s rumination circuit with real-time fMRI functional connectivity neurofeedback Hum Brain Mapp

67. Ramot M, Grossman S, Friedman D and Malach R 2016 Covert neurofeedback without awareness shapes cortical network spontaneous connectivity *Department of Neuroscience, Columbia University*, New York, NY 10027, USA. 113 E2413-20

68. Paret C, Goldway N, Zich C, Keynan J N, Hendler T, Linden D and Cohen Kadosh K 2019 Current progress in real-time functional magnetic resonance-based neurofeedback: Methodological challenges and achievements Neuroimage 202 116107

69. Sitaram R, Ros T, Stoeckel L, Haller S, Scharnowski F, Lewis-Peacock J, Weiskopf N, Blefari M L, Rana M, Oblak E, Birbaumer N and Sulzer J 2017 Closed-loop brain training: the science of neurofeedback Nat Rev Neurosci 18 86-100

70. Emmert K, Kopel R, Sulzer J, Brühl A B, Berman B D, Linden D E J, Horovitz S G, Breimhorst M, Caria A, Frank S, Johnston S, Long Z, Paret C, Robineau F, Veit R, Bartsch A, Beckmann C F, Van De Ville D and Haller S 2016 Meta-analysis of real-time fMRI neurofeedback studies using individual participant data: How is brain regulation mediated? NeuroImage 124, Part A 806-12

71. Mayeli A, Misaki M, Zotev V, Tsuchiyagaito A, Al Zoubi O, Phillips R, Smith J, Stewart J L, Refai H, Paulus M P and Bodurka J 2019 Self-regulation of ventromedial prefrontal cortex activation using real-time fMRI neurofeedback—Influence of default mode network Human Brain Mapping 41 342-52

