## Supplementary Materials for "The impact of real-time fMRI denoising on online evaluation of brain activity and functional connectivity"

#### fMRI samples for the simulation analyses

Resting-state fMRI data for eighty-seven healthy participants (45 females, mean age (SD) = 30 (10)) drawn from a previous study [1] was used for the simulation analyses. Magnetic resonance imaging was conducted on a whole-body 3 Tesla MR750 scanner (GE Healthcare, Milwaukee, WI). Single-shot gradient-recalled echo-planner imaging (EPI) with sensitivity encoding (SENSE) was used for fMRI. The scanning parameters were TR/TE = 2000/25 ms, FA = 75°, FOV = 240 mm, 34 axial slices with 2.9 mm thickness without gap, matrix = 96×96, SENSE acceleration factor R = 2. The EPI images were reconstructed into a 128×128 matrix resulting in 1.9×1.9×2.9 $mm^3$ voxel volume. The run time was 7 min 30 s (225 volumes). A photoplethysmograph with an infrared emitter placed under the pad of a participant's finger was used for pulse oximetry to measure cardiogram and a pneumatic respiration belt was used for respiration measurements. Physiological pulse oximetry and respiration waveforms were recorded with 40 Hz sampling frequency. The quality of the physiological signal recordings was checked visually. For anatomical reference, T1-weighted structural image was also acquired with SENSE accelerated magnetization-prepared rapid gradient-echo (MPRAGE) sequence (FOV = 240×192 mm, matrix = 256×256, 186 axial slices, slice thickness = 0.9 mm, 0.94×0.94×0.9 mm3 voxel volume, TR/TE = 5.0/2.0 ms, SENSE acceleration R = 2, flip angle = 8°, delay/inversion time = 1400/725 ms, sampling bandwidth = 31.2 kHz, scan time = 5 min 40 s).

Real-time fMRI neurofeedback task data for twenty-two participants (14 major depressive disorder (MDD) and 8 healthy control (HC) participants; 10 and 3 females in MDD and HC, respectively; mean age (SD) = 39 (9) and 34 (8) for MDD and HC, respectively) were drawn from the previous study [2]. Scanning parameters were the same as the resting-state data except TE = 30 ms, FA = 90°. Data at the first neurofeedback run was used for the simulation. In the neurofeedback run, the 40s-block of rest, happy memory recall with the left amygdala neurofeedback (LA-NF), and the count task were repeated four times. During the neurofeedback block, participants were required to increase the LA-NF signal show on the screen by a red vertical bar by recalling a happy autobiographical memory. The scan time was 8m46s with additional 6s rest in the first and one 40s rest block at the last. The details of the experimental procedures were described in the previous report [2]. Physiological signals and T1-weighted structural image were also acquired with the procedures in the resting-state.

#### RTP simulation system implementation

The RTP simulation system was built based on our previous implementation [3]. While the previous system was implemented as an AFNI (https://afni.nimh.nih.gov/) plugin with C language, the present system was implemented with python3, independent of AFNI, for portability and utility of the simulation analysis. The code for the simulated system is available on the GitHub site (https://github.com/mamisaki/fMRI_RTP_Simulation). The system was composed of five processing modules; WATCH, TSHIFT, VOLREG, SMOOTH, and REGRESS.

The WATCH module monitors a directory where a new fMRI volume file is created in real-time and loads the file in order to send the data to the processing modules.

The TSHIFT module adjusts timing differences between slices by interpolating and resampling the time-course data at a shifted time point. We note that while Heunis, Lamerichs, Zinger, Caballero-Gaudes, Jansen, Aldenkamp and Breeuwer [4] described that there is no algorithmic difference between the offline and real-time slice-timing corrections, they could be different depending on the order of temporal interpolation [3]. Since a future data point is not available in RTP, a high-order interpolation cannot be used as it requires multiple future time points. The data were resampled at the earliest slice timing in a volume to avoid extrapolation. The system has linear and cubic interpolation options for resampling the data [3]. The present simulation used the cubic interpolation because a previous report [3] demonstrated that the cubic interpolation adapted to RTP had enough accuracy, similar to an offline one. The TSHIFT process started after receiving three volumes (except for the initial three volumes, which were discarded to ensure a steady-state fMRI signal) in order to wait for getting enough samples for the interpolation. The volumes before starting the process (after the steady state) were also applied the correction retrospectively and sent to the next process.

The VOLREG module performs volume registration for motion correction. We used the AFNI *3dvolreg* function by compiling its source code as a C-shared library. The RTP system called it via python's *ctypes* interface. The present simulation used the cubic interpolation for resampling a volume at a motion-corrected space. The reference volume for the registration was taken from another functional scan in the same session with the same imaging resolution.

The SMOOTH module performs spatial smoothing by convolving a Gaussian kernel. We used AFNI implementation of *3dBlurInMask* by compiling it in a C-shared library. The smoothing was applied within the brain mask. Smoothing with 6mm-FWHM Gaussian was used in the simulation.

The REGRESS module performs signal scaling to a percent change in each voxel and then regression with multiple regressors. The regressors can include Legendre polynomials for high-pass filtering, motion parameters, motion derivatives, global signal, mean signals of white matter and ventricle region, RETROICOR [5], and RVT [6]. The polynomial order was determined by the length of the scan at the time of the regression, calculated as 1 + int($d$/150), where $d$ is the scan duration in seconds. This order is the default in AFNI's *3dDecovlove*. The global signal and the white matter and ventricle average signals were calculated with unsmoothed data, the VOLREG module's output.

Calculations of RETROICOR and RVT models were implemented in a C library function based on the AFNI RetroTS.m MATLAB script. The calculation was performed at every TR using the accumulated history of the cardiogram and respiration signals from the start of the scan until the current TR. We used the AFNI's default implementation and no specific adaptation for a real-time application was made. The only difference between real-time and offline processes was that the signals until the present TR were used in a real-time process while the signals of whole time-series in a run were used in the offline analysis. Eight RETROICOR (four Fourierbasis functions for cardiac and respiration, respectively), and five RVT regressors were created in the simulation.

The system implemented the cumulative GLM for regression analysis with an ordinary least square (OLS) approach. We used the PyTorch library (https://pytorch.org/) to solve the OLS on GPU or CPU. The beta values were used to regress out noise components to get the residual as a denoised signal [7]. With GPU implementation, data transfer from system memory to the GPU device could be a bottleneck for the processing time. This overhead was minimized by keeping memory on GPU at the beginning large enough for the expected maximum length of the scan volumes. Then, only new data received in real-time with an applied mask was sent to the pre-assigned GPU memory space. The REGRESS process began after receiving the number of volumes equal to the number of regressors plus one. The mean signal for the scaling was calculated with the volumes before starting the GLM analysis [3].

Motion parameters were received from the VOLREG module. The white matter and ventricle masks were defined in the MNI template brain, warped into the individual anatomy image aligned to the VOLREG reference image, and then resampled into the function image resolution. The masks were eroded two and one voxels for white matter and ventricle masks, respectively, in the individual anatomical image resolution to avoid including gray matter signals. The masks were made in offline before the RTP simulation. We used *align_epi_anat.py* in AFNI to align an anatomy image to a function image and the Advanced Normalization Tools (ANTs, http://stnava.github.io/ANTs/) Avants, Epstein, Grossman and Gee [8] to warp the MNI template to an individual anatomy image. The global signal regressor was the mean signal of the whole-brain region, which was the intersection of the anatomical brain mask and the signal mask made with 3dAutomask in AFNI.

#### Offline fMRI processing

The offline fMRI processing was performed with AFNI. The process included despike, RETROICOR and RVT regression, slice-timing and motion corrections, smoothing with 6mm-FWHM kernel, scaling to percent change relative to the mean signal in each voxel and regression with 12 motion parameters (three shifts and three rotations and their temporal derivatives), three principal components of ventricle signals, local white matter average signals (ANATICOR [9]), and Legendre polynomials for high-pass filtering. We used the default parameters of AFNI in offline processing.

#### Table S1.

Statistical values of the RTP noise reduction performance for voxel-wise signals. *P*-values are corrected for multiple comparisons.

| | Noise | | | | | |
|---|---|---|---|---|---|---|
| | Motion | | Cardiac | | Respiration | |
| | *t* | *p* | *t* | *p* | *t* | *p* |
| RTP1 – RTP0 (+TSHIFT) | 1.395 | 0.791 | 0.001 | 1.000 | -1.229 | 0.883 |
| RTP2 – RTP1 (+SMOOTH) | 1.692 | 0.577 | **3.871** | **0.001** | **3.498** | **0.005** |
| RTP3 – RTP2 (+REG[HPF]) | **4.327** | **< 0.001** | **7.978** | **< 0.001** | **11.099** | **< 0.001** |
| RTP4 – RTP3 (+REG[Mot]) | **-8.782** | **< 0.001** | 0.377 | 1.000 | **-9.063** | **< 0.001** |
| RTP5 – RTP4 (+REG[dMot]) | -2.662 | 0.076 | -2.469 | 0.127 | **-3.587** | **0.004** |
| RTP6 – RTP5 (+REG[GS]) | 0.134 | 1.000 | **3.373** | **0.008** | -0.348 | 1.000 |
| RTP7 – RTP6 (+REG[WM, Vent]) | -0.421 | 1.000 | **-3.921** | **0.001** | -1.071 | 0.944 |
| RTP8 – RTP7 (+REG[RETROICOR]) | -0.258 | 1.000 | **-21.449** | **< 0.001** | **-3.919** | **0.001** |
| RTP9 – RTP8 (+REG[RVT]) | -0.241 | 1.000 | 0.181 | 1.000 | 1.084 | 0.940 |
| Offline – RTP9 | -0.317 | 1.000 | **-6.417** | **< 0.001** | **-14.098** | **< 0.001** |

#### Table S2.

Statistical values of the RTP noise reduction performance for the 5-TR sliding-window connectivity time-course. *P*-values are corrected for multiple comparisons.

| | Noise | | | | | |
|---|---|---|---|---|---|---|
| | Motion | | Cardiac | | Respiration | |
| | *t* | *p* | *t* | *p* | *t* | *p* |
| RTP1 – RTP0 (+TSHIFT) | 0.014 | 1.000 | -1.658 | 0.603 | -0.174 | 1.000 |
| RTP2 – RTP1 (+SMOOTH) | 0.688 | 0.997 | 1.518 | 0.707 | 0.966 | 0.970 |
| RTP3 – RTP2 (+REG[HPF]) | -0.541 | 1.000 | -0.218 | 1.000 | -1.159 | 0.913 |
| RTP4 – RTP3 (+REG[Mot]) | 0.396 | 1.000 | -1.855 | 0.454 | -2.184 | 0.247 |
| RTP5 – RTP4 (+REG[dMot]) | **-3.545** | **0.004** | -1.623 | 0.629 | **-3.633** | **0.003** |
| RTP6 – RTP5 (+REG[GS]) | **-3.143** | **0.018** | -0.572 | 0.999 | -0.802 | 0.991 |
| RTP7 – RTP6 (+REG[WM, Vent]) | -0.393 | 1.000 | -1.740 | 0.541 | -0.372 | 1.000 |
| RTP8 – RTP7 (+REG[RETROICOR]) | -0.460 | 1.000 | **-3.938** | **0.001** | -0.228 | 1.000 |
| RTP9 – RTP8 (+REG[RVT]) | 0.586 | 0.999 | 0.895 | 0.981 | 1.732 | 0.547 |
| Offline – RTP9 | -0.269 | 1.000 | -0.429 | 1.000 | -1.966 | 0.377 |

**Table S3.**

Statistical values of the RTP noise reduction performance for the two-point connectivity time-course. P-values are corrected for multiple comparisons.

| | Noise | | | | | |
|---|---|---|---|---|---|---|
| | Motion | | Cardiac | | Respiration | |
| | *t* | *p* | *t* | *p* | *t* | *p* |
| RTP1 – RTP0 (+TSHIFT) | 1.232 | 0.882 | **-3.140** | **0.018** | **-5.065** | **< 0.001** |
| RTP2 – RTP1 (+SMOOTH) | 1.185 | 0.903 | **3.695** | **0.003** | 0.569 | 0.999 |
| RTP3 – RTP2 (+REG[HPF]) | -0.541 | 1.000 | -0.523 | 1.000 | -0.332 | 1.000 |
| RTP4 – RTP3 (+REG[Mot]) | -0.404 | 1.000 | **-2.928** | **0.035** | -1.705 | 0.567 |
| RTP5 – RTP4 (+REG[dMot]) | -0.895 | 0.981 | **-3.243** | **0.013** | **-3.752** | **0.002** |
| RTP6 – RTP5 (+REG[GS]) | -2.376 | 0.160 | 2.211 | 0.232 | 0.364 | 1.000 |
| RTP7 – RTP6 (+REG[WM, Vent]) | -0.200 | 1.000 | **-3.029** | **0.026** | -0.776 | 0.993 |
| RTP8 – RTP7 (+REG[RETROICOR]) | -0.380 | 1.000 | **-8.464** | **< 0.001** | -0.950 | 0.972 |
| RTP9 – RTP8 (+REG[RVT]) | 0.534 | 1.000 | 0.271 | 1.000 | 2.071 | 0.310 |
| Offline – RTP9 | -1.700 | 0.571 | -1.610 | 0.639 | **-3.619** | **0.003** |

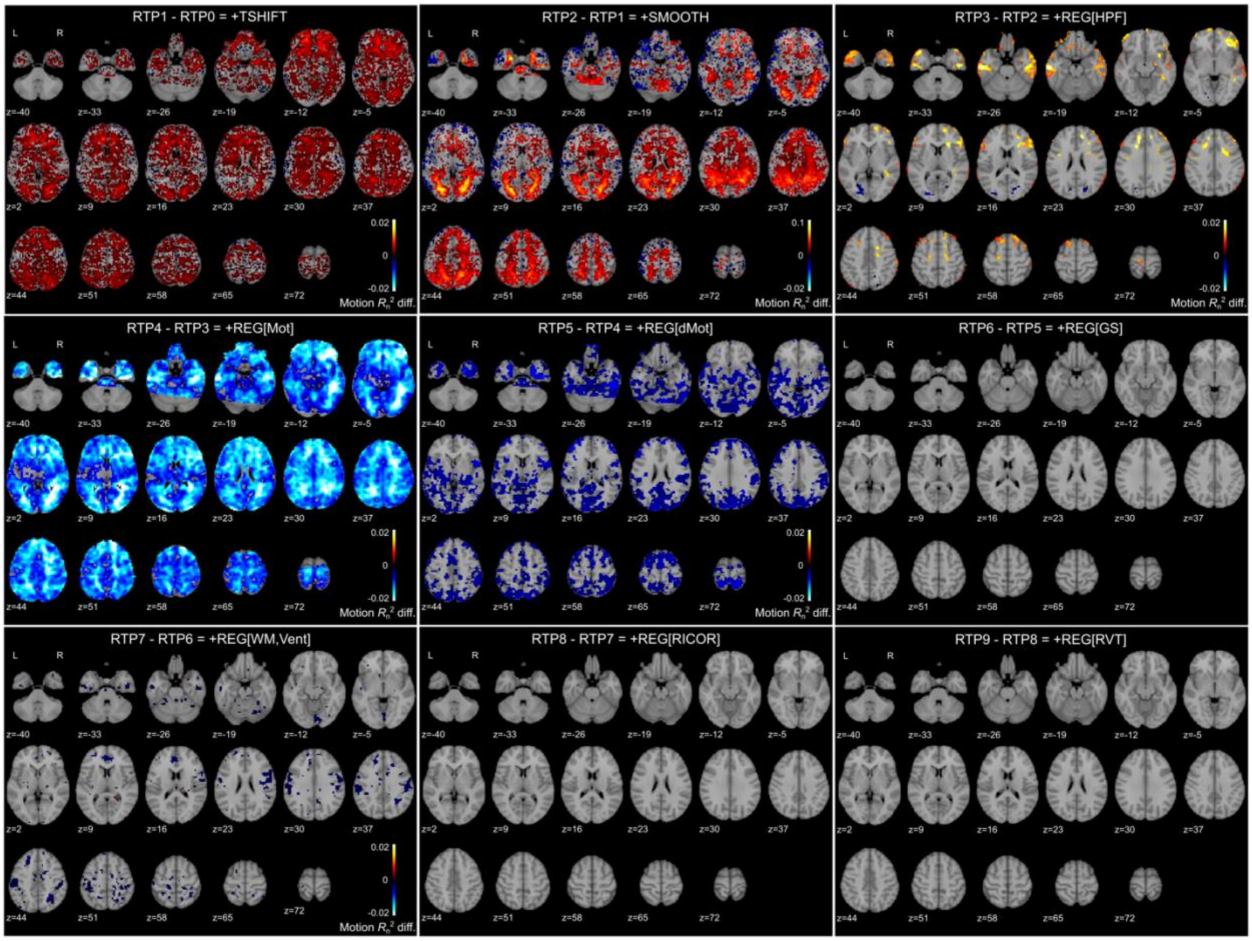

**Figure S1.**

Significant $R_n^2$ difference of motion noise in the voxel-wise signal between real-time processing pipelines. The maps were thresholded by FDR < 0.05 with a randomization test.

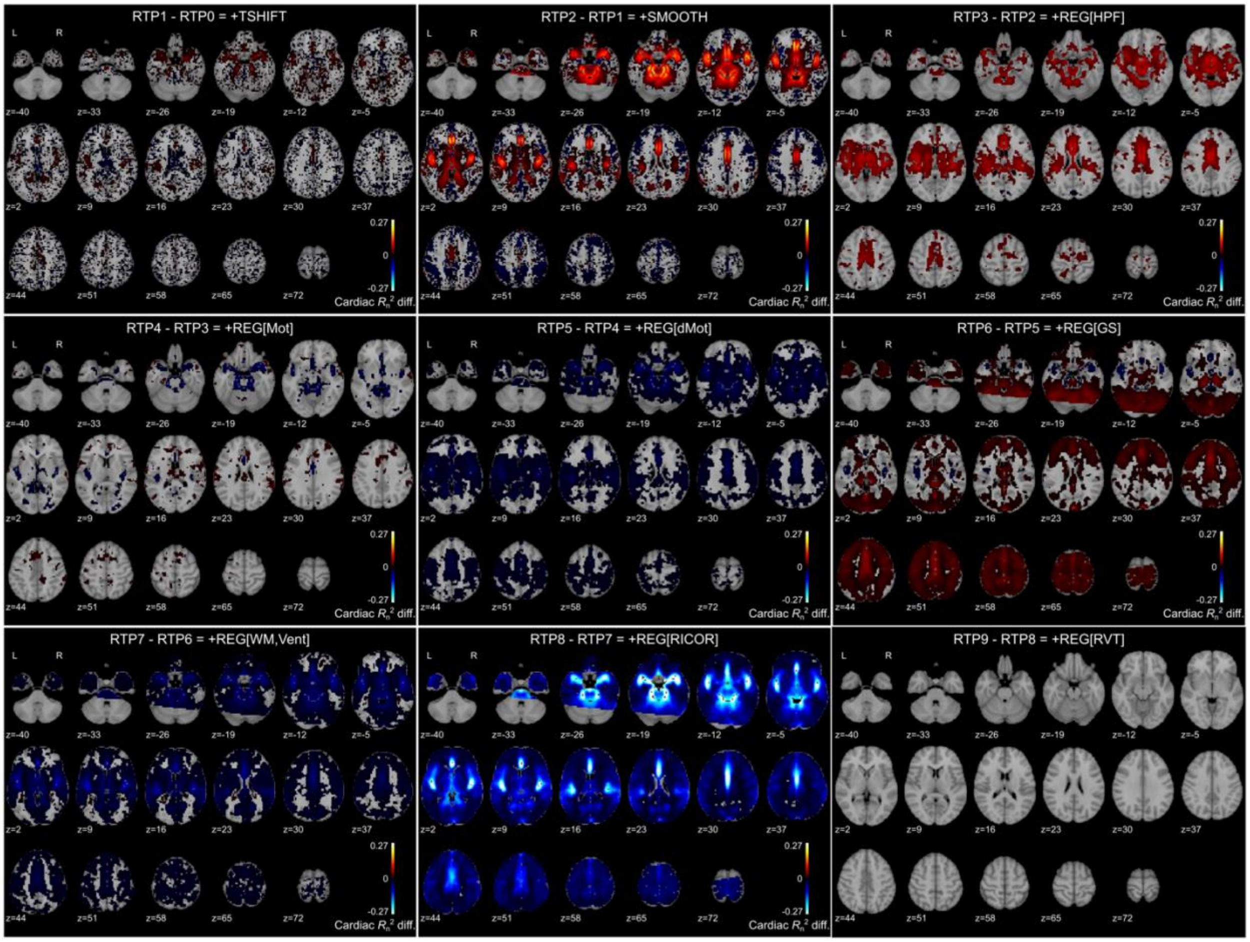

#### Figure S2.

Significant $R_n^2$ difference of cardiac noise in the voxel-wise signal between real-time processing pipelines. The maps were thresholded by FDR < 0.05 with a randomization test.

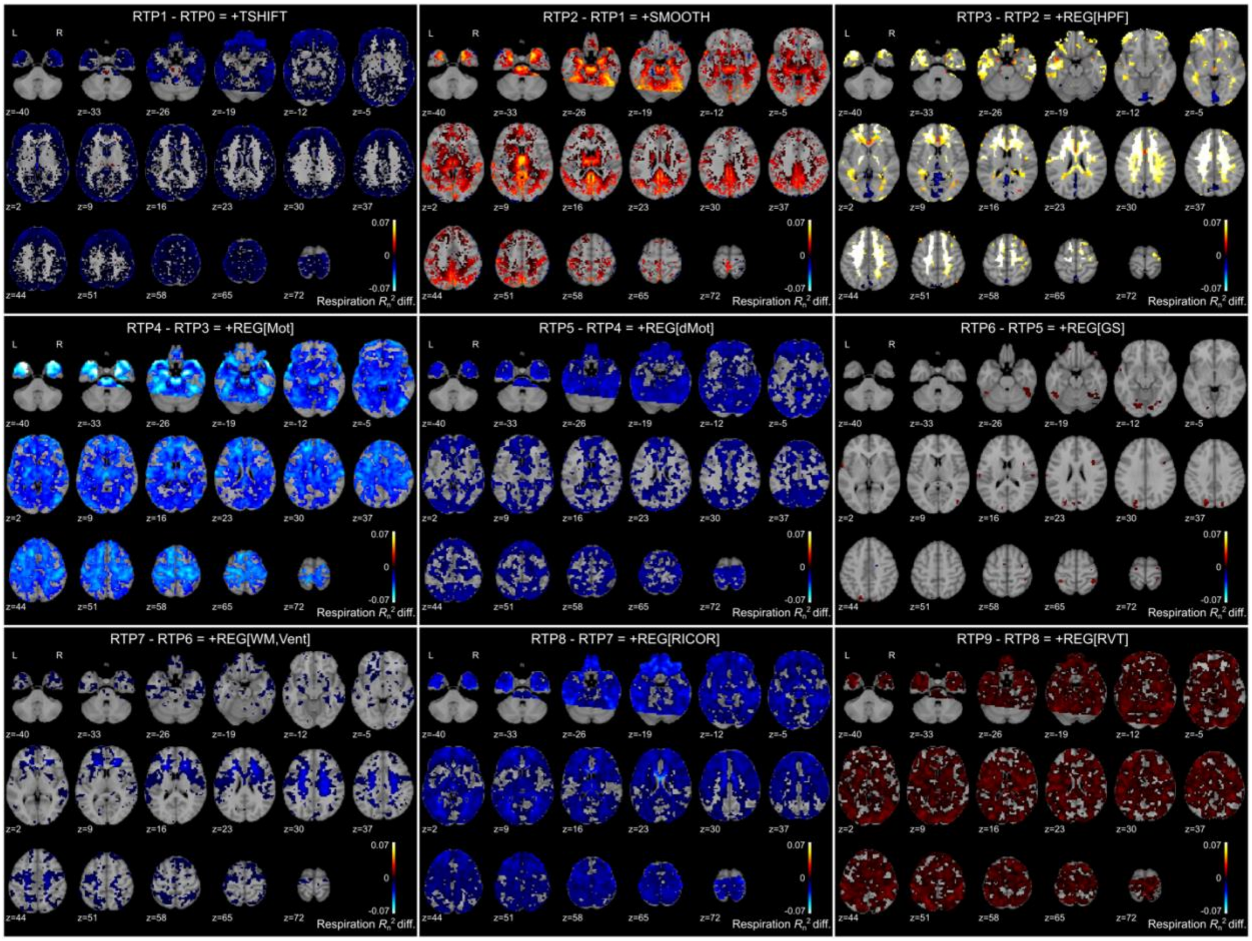

**Figure S3.**

Significant $R_n^2$ difference of respiration noise in the voxel-wise signal between real-time processing pipelines. The maps were thresholded by FDR < 0.05 with a randomization test.

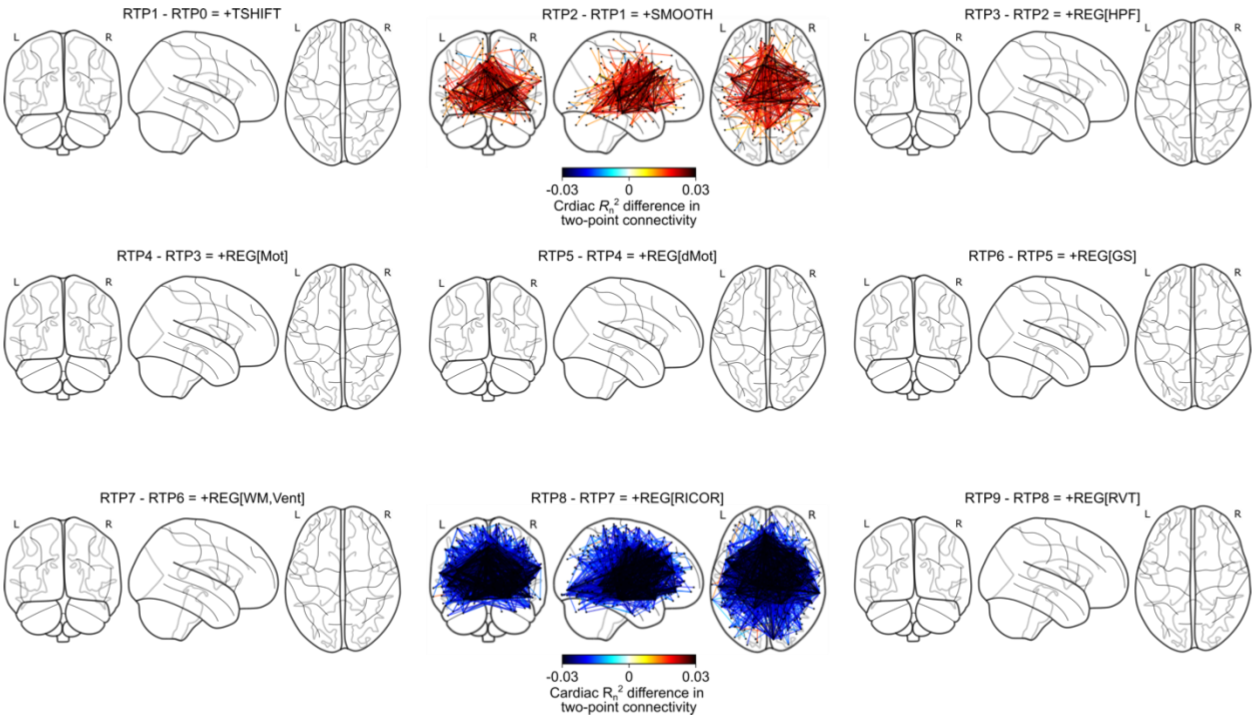

**Figure S4.**

Connectivity plots of significant difference of $R_n^2$ of cardiac noise in two-point connectivity time-course between real-time processing pipelines. The plots were thresholded by FDR < 0.05 with randomization test.

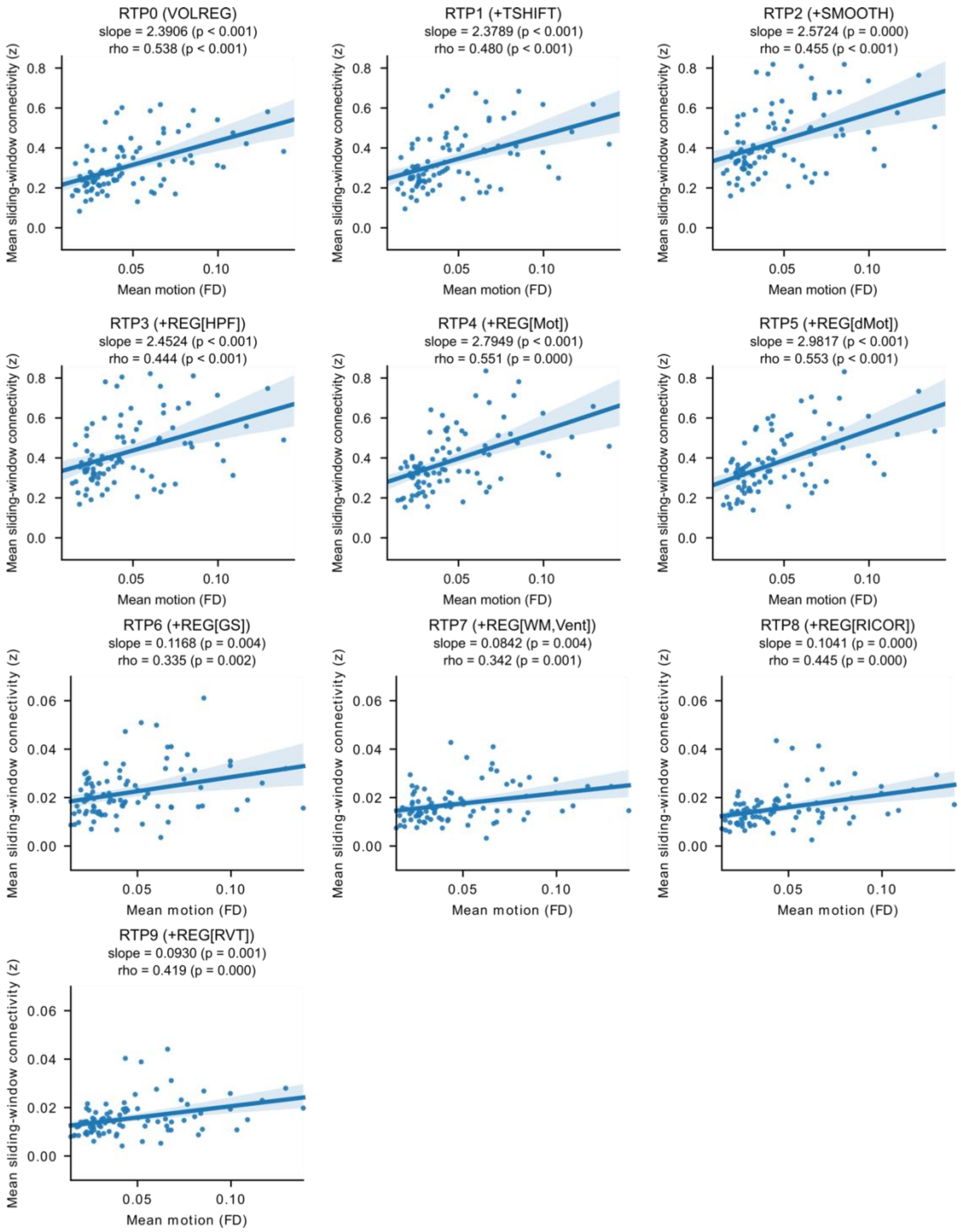

**Figure S5.**

Association between the mean sliding-window (5-TR width) connectivity and the mean motion (frame-wise displacement, FD). Each point indicates a participant. The shadow around the line indicates a 95% confidence interval. 'Slope' is a fitted coefficient of the motion in linear regression analysis. 'Rho' is the Spearman's rank-order correlation.

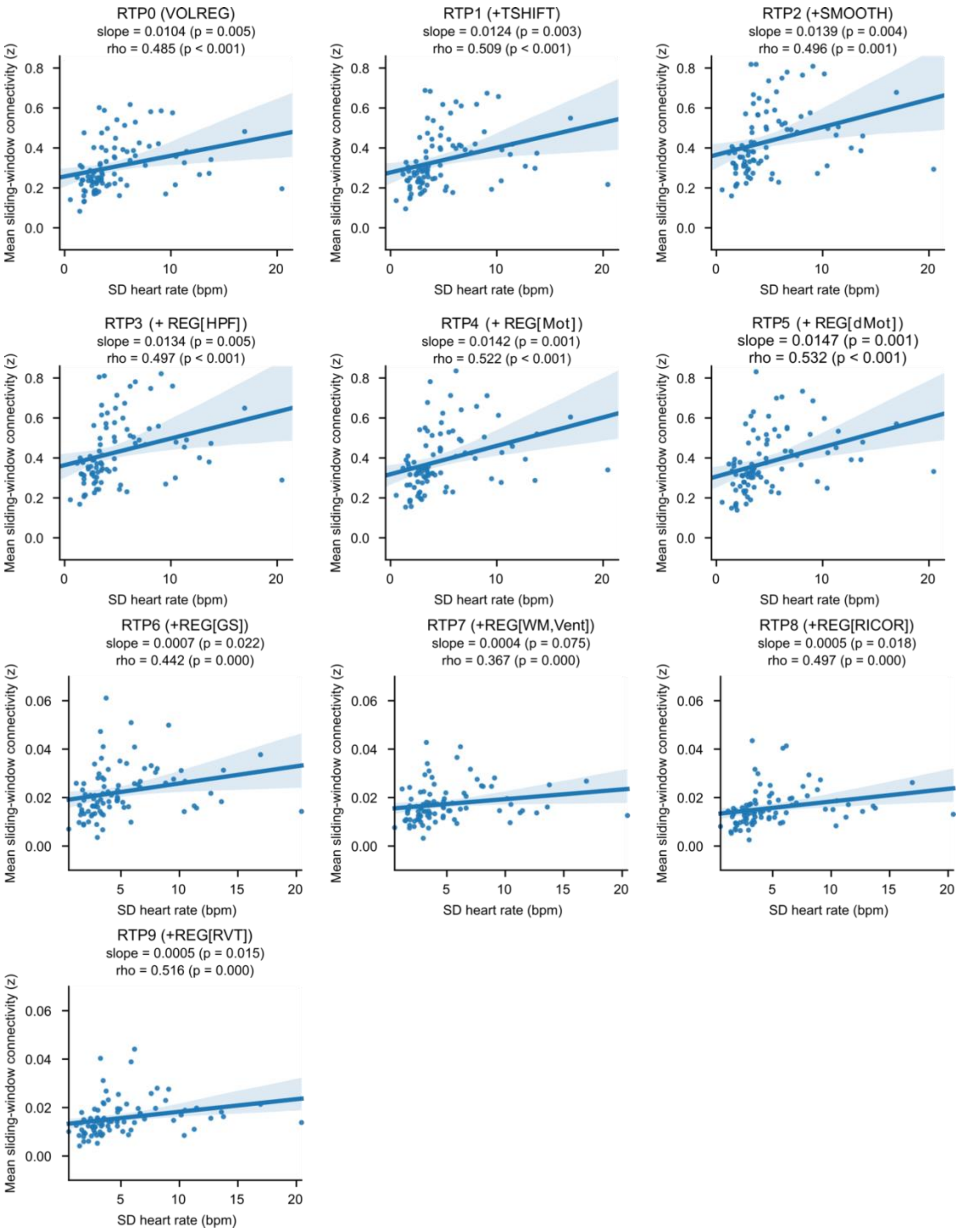

**Figure S6.**

Association between the mean sliding-window (5-TR) connectivity and the standard deviation (SD) of heart rate. Each point indicates a participant. The shadow around the line indicates a 95% confidence interval. 'Slope' is a fitted coefficient of the SD heart rate in linear regression analysis. 'Rho' is Spearman's rank-order correlation.

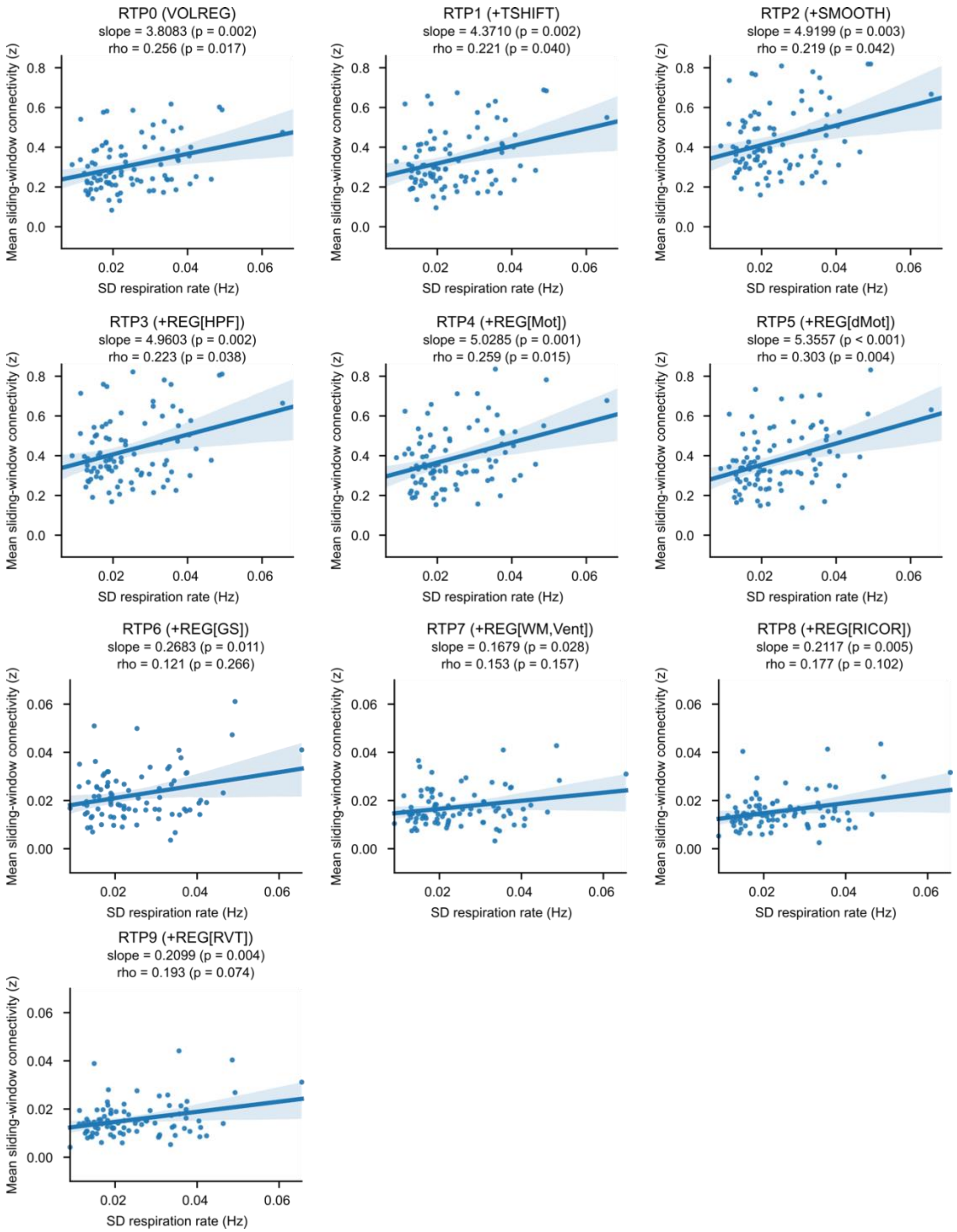

**Figure S7.**

Association between the mean sliding-window (5-TR) connectivity and the standard deviation (SD) of respiration rate. Each point indicates a participant. The shadow around the line indicates a 95% confidence interval. 'Slope' is a fitted coefficient of the SD heart rate in linear regression analysis. 'Rho' is Spearman's rank-order correlation.

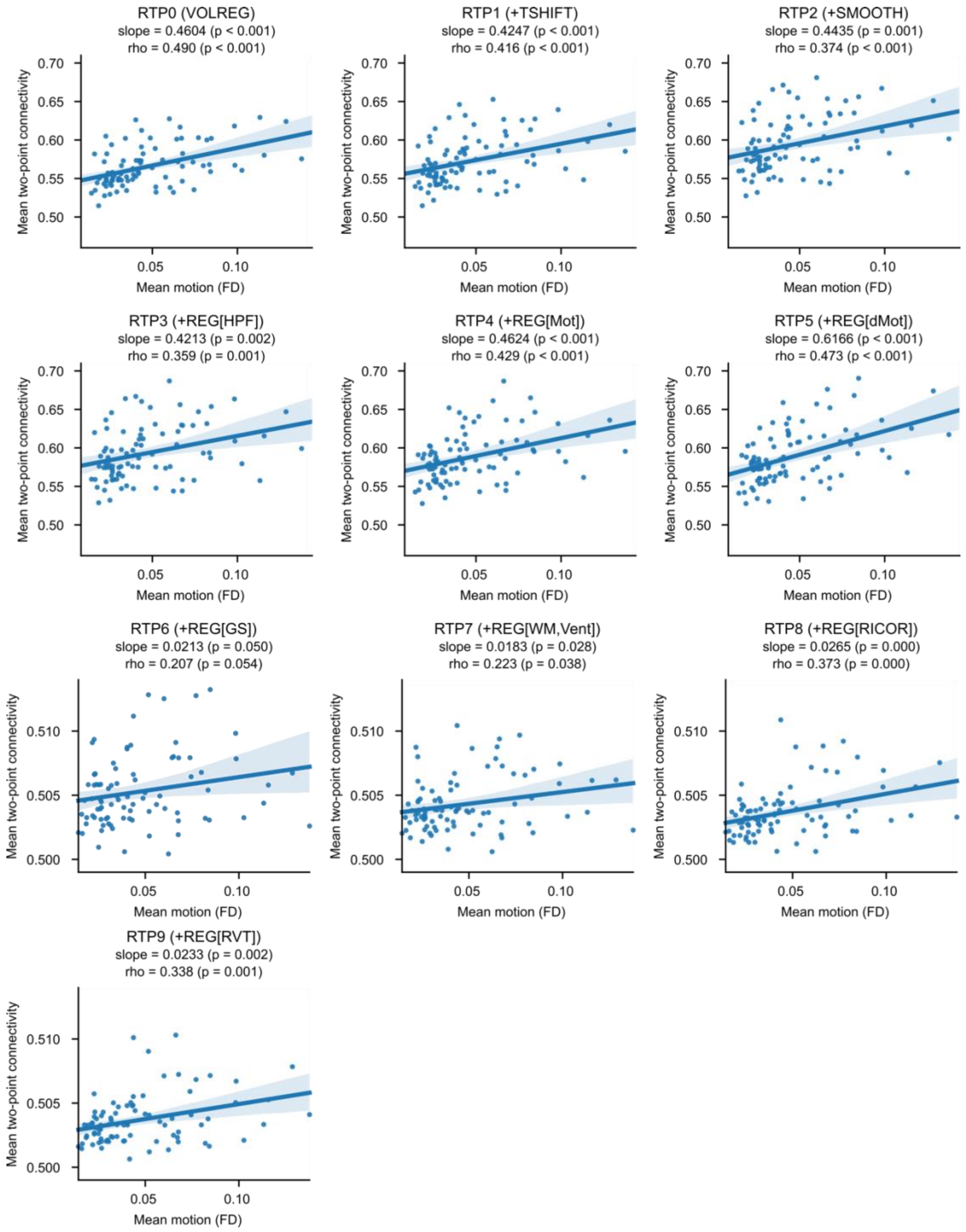

**Figure S8.**

Association between the mean two-point connectivity and the mean motion (frame-wise displacement, FD). Each point indicates a participant. The shadow around the line indicates a 95% confidence interval. 'Slope' is a fitted coefficient of the motion in linear regression analysis. 'Rho' is Spearman's rank-order correlation.

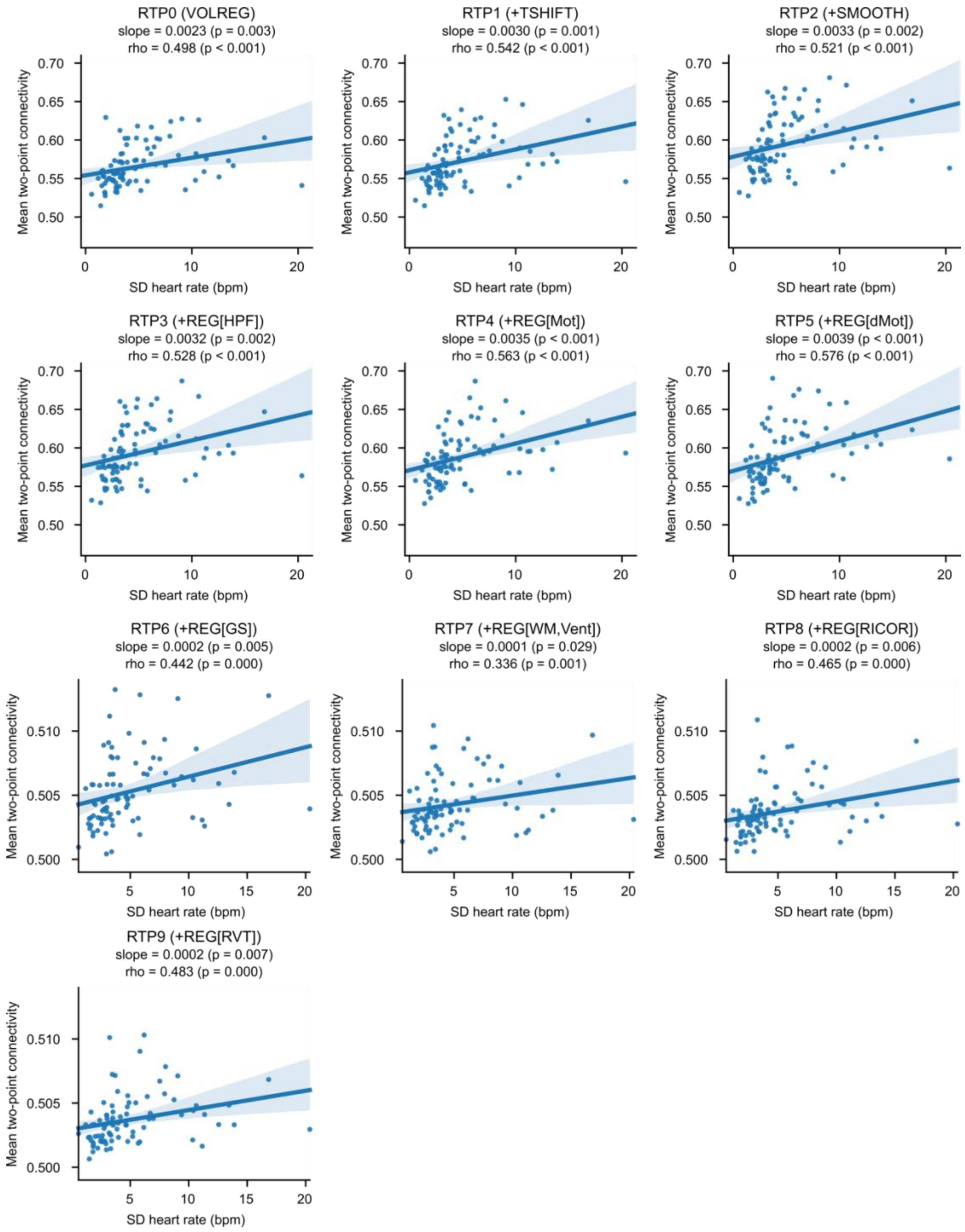

#### Figure S9.

Association between the mean two-point connectivity and the standard deviation (SD) of heart rate. Each point indicates a participant. The shadow around the line indicates a 95% confidence interval. 'Slope' is a fitted coefficient of the SD heart rate in linear regression analysis. 'Rho' is Spearman's rank-order correlation.

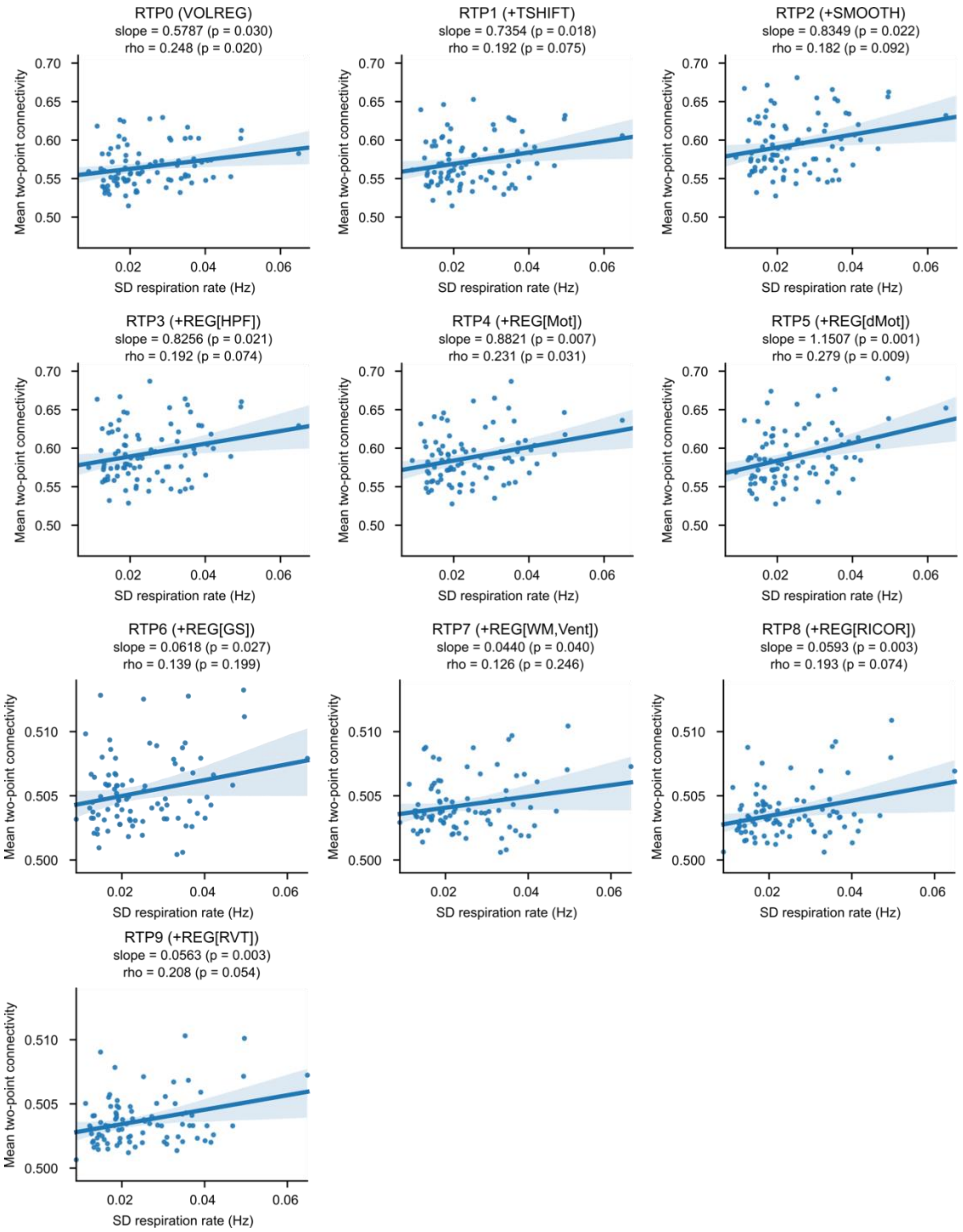

**Figure S10.**

Association between the mean two-point connectivity and the standard deviation (SD) of respiration rate. Each point indicates a participant. The shadow around the line indicates a 95% confidence interval. 'Slope' is a fitted coefficient of the SD heart rate in linear regression analysis. 'Rho' is Spearman's rank-order correlation.

### References

[1] Misaki M, Suzuki H, Savitz J, Drevets W C and Bodurka J 2016 Individual Variations in Nucleus Accumbens Responses Associated with Major Depressive Disorder Symptoms *Scientific reports* **6** 21227

[2] Young K D, Zotev V, Phillips R, Misaki M, Yuan H, Drevets W C and Bodurka J 2014 Real-time FMRI neurofeedback training of amygdala activity in patients with major depressive disorder *PLoS One* **9** e88785

[3] Misaki M, Barzigar N, Zotev V, Phillips R, Cheng S and Bodurka J 2015 Real-time fMRI processing with physiological noise correction – Comparison with off-line analysis *Journal of Neuroscience Methods* **256** 117-21

[4] Heunis S, Lamerichs R, Zinger S, Caballero-Gaudes C, Jansen J F A, Aldenkamp B and Breeuwer M 2020 Quality and denoising in real-time functional magnetic resonance imaging neurofeedback: A methods review *Hum Brain Mapp* **41** 3439-67

[5] Glover G H, Li T Q and Ress D 2000 Image-based method for retrospective correction of physiological motion effects in fMRI: RETROICOR *Magn Reson Med* **44** 162-7

[6] Birn R M, Smith M A, Jones T B and Bandettini P A 2008 The respiration response function: the temporal dynamics of fMRI signal fluctuations related to changes in respiration *NeuroImage* **40** 644-54

[7] Hinds O, Ghosh S, Thompson T W, Yoo J J, Whitfield-Gabrieli S, Triantafyllou C and Gabrieli J D E 2011 Computing moment-to-moment BOLD activation for real-time neurofeedback *NeuroImage* **54** 361-8

[8] Avants B B, Epstein C L, Grossman M and Gee J C 2008 Symmetric diffeomorphic image registration with cross-correlation: evaluating automated labeling of elderly and neurodegenerative brain *Med. Image Anal* **12** 26-41

[9] Jo H J, Saad Z S, Simmons W K, Milbury L A and Cox R W 2010 Mapping sources of correlation in resting state FMRI, with artifact detection and removal *NeuroImage* **52** 571-82
